# Quinolinic acid links kidney injury to brain toxicity

**DOI:** 10.1101/2024.05.07.592801

**Authors:** Afaf Saliba, Subrata Debnath, Ian Tamayo, Jana Tumova, Meyer Maddox, Pragya Singh, Caitlyn Fastenau, Soumya Maity, Hak Joo Lee, Guanshi Zhang, Leila Hejazi, Jason C. O’Connor, Bernard Fongang, Sarah C Hopp, Kevin F. Bieniek, James D. Lechleiter, Kumar Sharma

## Abstract

Kidney dysfunction often leads to neurological impairment, yet the complex kidney-brain relationship remains elusive. We employed spatial and bulk metabolomics to investigate a mouse model of rapid kidney failure induced by mouse double minute 2 (*Mdm2)* conditional deletion in the kidney tubules to interrogate kidney and brain metabolism. Pathway enrichment analysis of focused plasma metabolomics panel pinpointed tryptophan metabolism as the most altered pathway with kidney failure. Spatial metabolomics showed toxic tryptophan metabolites in the kidneys and brains, revealing a novel connection between advanced kidney disease and accelerated kynurenine degradation. In particular, the excitotoxic metabolite quinolinic acid was localized in ependymal cells adjacent to the ventricle in the setting of kidney failure. These findings were associated with brain inflammation and cell death. A separate mouse model of acute kidney injury also had an increase in circulating toxic tryptophan metabolites along with altered brain inflammation. Patients with advanced CKD similarly demonstrated elevated plasma kynurenine metabolites and quinolinic acid was uniquely correlated with fatigue and reduced quality of life in humans. Overall, our study identifies the kynurenine pathway as a bridge between kidney decline, systemic inflammation, and brain toxicity, offering potential avenues for diagnosis and treatment of neurological issues in kidney disease.

## INTRODUCTION

Acute and chronic kidney disease (CKD), characterized by decline in kidney function, are global health burdens and can have a major impact on neurologic dysfunction (1). In the USA, acute kidney injury (AKI) has a prevalence of 7% in hospitalized patients and is associated with increased mortality risk (2, 3), and CKD has a prevalence of 14% (4, 5) and also associated with increased mortality (6–8). AKI and CKD patients are more prone to complications, including cardiovascular, metabolic, neurologic, and other disturbances (5). Fatigue and depression are common neurolopsychiatric syndromes experienced by CKD patients as kidney function progresses to stage 4 or 5 (9) however the basis for these symptoms are unclear.

Kidney-related neurologic dysfunction has been associated with ischemic cerebrovascular lesions (10), white matter lesions (11), microbleeds (12), and circulating uremic toxins (13, 14). However, the underlying mechanisms affecting the kidney-brain axis remain poorly understood. To interrogate how reduced kidney function may lead to neurologic dysfunction, we analyzed candidate molecules via a targeted metabolomics analysis of 32 metabolites in two separate mouse models of rapid kidney decline followed by spatial metabolomics in kidney and brain. In addition, clinical translation studies were performed in patients with stage 4 and stage 5 CKD. We found accumulation of circulating toxic metabolites produced by the catabolism of essential amino acid tryptophan (Trp) via the kynurenine pathway (KP) in the plasma, kidney, and brain. Clinical samples demonstrated accumulation of the same neurotoxic metabolites in plasma samples in patients with advanced CKD. In both pre-clinical and clinical studies, the neurotoxic metabolite quinolinic acid emerged as a key link in the kidney-brain axis.

## RESULTS

### Tryptophan metabolism is altered in plasma of mice with severe kidney dysfunction

Based on previous studies highlighting the role of mouse double minute 2 (*Mdm2)* in human and experimental kidney disease (15, 16) we studied rapid kidney failure in the doxycycline (Dox)-induced conditional knockout of *Mdm2* targeting renal tubular epithelial cells (*Mdm2*cKO). As depicted in (**Supplemental Figure 1**), mice with inducible kidney tubular Mdm2 deficiency exhibit pronounced kidney dysfunction within a span of 3-7 days. This dysfunction was marked by severe tubular cell damage and a marked decline in kidney function, evidenced by elevated blood urea nitrogen (BUN) and plasma creatinine levels.

We performed targeted bulk metabolomics in plasma samples from *Mdm2*cKO (n=12) vs control (n=9) mice, at day 6 of Dox administration, with a panel of 32 amino acid-related metabolites relevant to human kidney disease, based on a prior untargeted metabolomic analysis from over 1000 patients (17). Unbiased pathway enrichment analyses of the targeted panel revealed Trp metabolism as the top enriched pathway with the lowest false discovery rate (FDR) value (p<0.001) (**Figure 1A**). Trp pathway metabolites in the plasma of *Mdm2*cKO mice showed a significant decrease in Trp and serotonin (5-HT) and an increase in kynurenine (KYN), 3-hydroxykynurenine (3HK), and quinolinic acid (QA) (**Figure 1, B-F**). These changes suggest a shift in Trp metabolism away from 5-HT production towards the kynurenine pathway (KP). KP is typically activated by enzymes such as indoleamine 2,3-dioxygenase (IDO) and kynurenine 3-monooxygenase (KMO), which are often upregulated in response to inflammatory cytokines. Interestingly, the plasma KYN-to-Trp ratio, an indicator of IDO activity, and the plasma 3HK-to-KYN ratio, an indicator of KMO activity, were significantly correlated with plasma creatinine levels from day 3 to day 6 of Dox administration to the *Mdm2*cKO mice (r=0.8562; p<0.0001; r=0.8527; p<0.0001 respectively) (**Supplemental Figure 1, E-F and Supplemental Figure 2**), indicating a strong link between altered tryptophan metabolism and kidney function.

**Figure 1.**
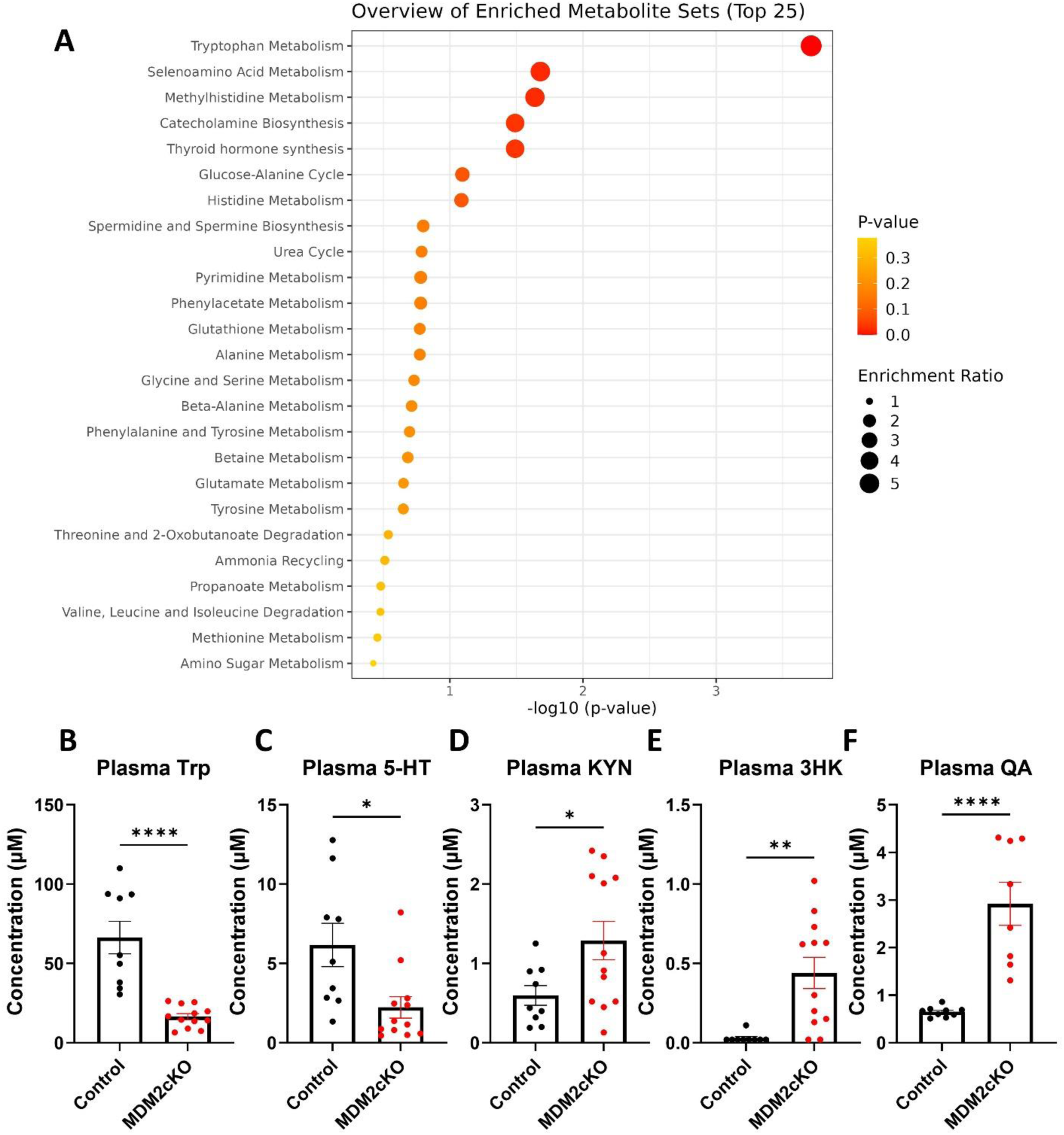
Plasma metabolomics highlight tryptophan metabolism as the most significantly skewed pathway in mice with kidney dysfunction. Targeted metabolomics measured via CE-MS ZipChip, incorporating 32 metabolites was performed in plasma from *Pax8*-rtTAcre;*Mdm2*f/f (n=12; *Mdm2* Conditional Knockout-*Mdm2*cKO) mice and *Mdm2*f/f mice (n=9; control), at day 6 of doxycycline (Dox). (A) Overview of unbiased pathway enrichment analysis on MetaboAnalyst 5.0 (library: SMPDB). Raw tryptophan metabolite concentrations are displayed: (B) tryptophan (Trp) (C) serotonin (5-HT) (D) kynurenine (KYN) (E) 3-hydroxykynurenine (3HK) (F) quinolinic acid (QA). Graphs display means ± SEM. Two-tailed t-tests with significance levels set at *p < 0.05, **p < 0.01, and ****p < 0.0001.

### Kidney failure is associated with increased inflammation and tryptophan pathway dysregulation in mice

*Mdm2*cKO mice also exhibited an increase in transformation related protein 53 (*Trp53*) mRNA levels, alongside the downstream target of p53, cyclin dependent kinase inhibitor 1A (*Cdkn1a*), indicating a response to cell stress. In addition, there was an increase in hypoxia inducible factor 1, alpha subunit (*Hif1a*), and NADPH oxidase 2 (*Nox2*) mRNA levels indicating stimulation of hypoxic and oxidative stress with kidney failure (**Figure 2, A-D)**. Several inflammatory markers were also upregulated in the kidney (**Figure 2, E-H)**. These include C-C motif chemokine ligand 2 (*Ccl2*), which recruits monocytes, memory T cells, and dendritic cells to sites of tissue injury; C-X-C motif chemokine Ligand 1 (*Cxcl1*) which mediates migration of immune cells to the inflammation site; and interleukin 6 (*Il6*), a central cytokine in inflammation along with interleukin 1 beta (*Il1b*).

**Figure 2.**
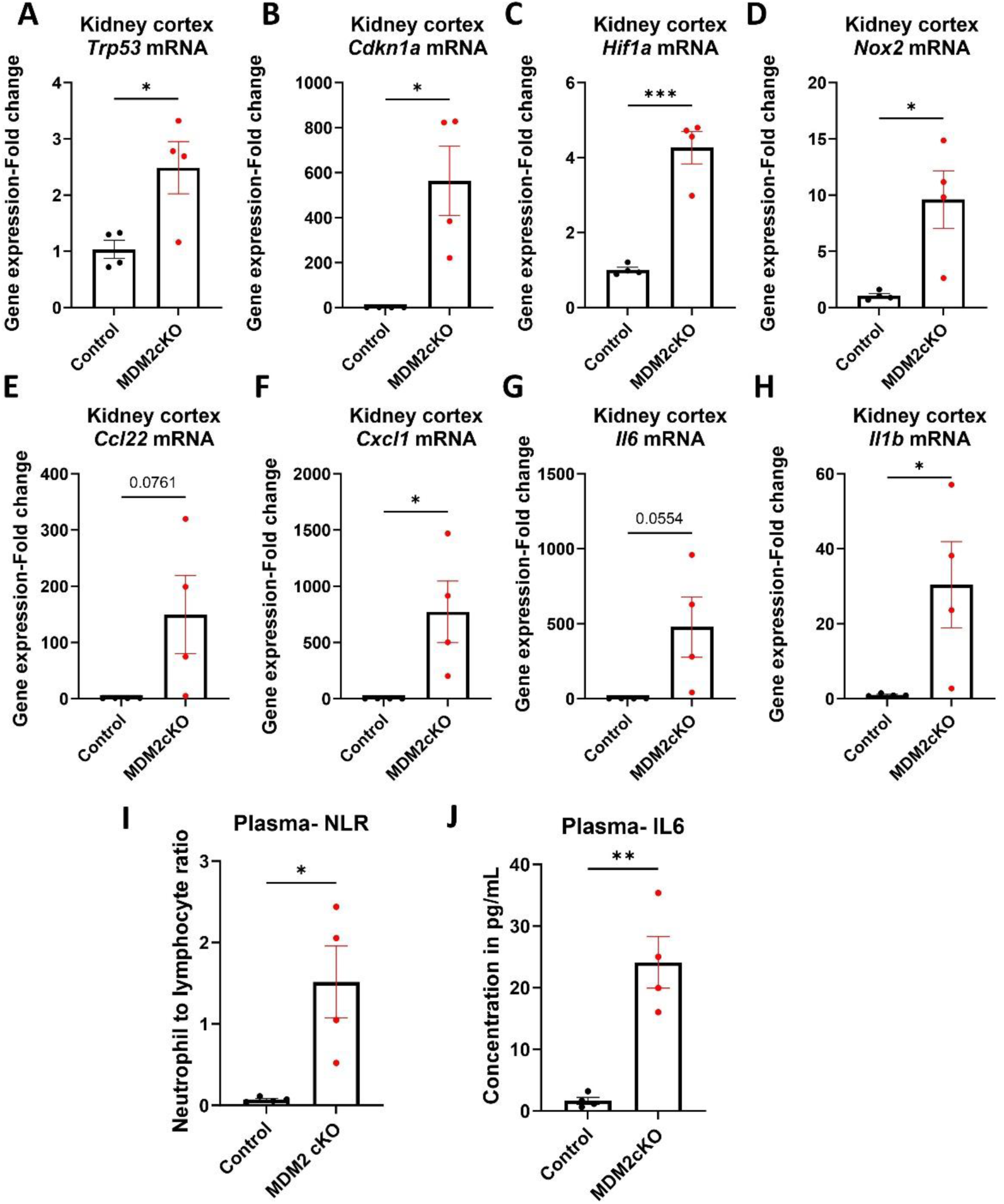
Tubular kidney cell death is associated with increased inflammation locally at the kidney cortex and at the systemic level. Kidney cortex RNA lysates were prepared from *Mdm2*cKO (n=4) vs. control mice (n=4) at 6 days of doxycycline (Dox) administration in drinking water 2mg/ml. All qPCR results are normalized to GAPDH expression, displayed here in fold change vs. control: (A) transformation related protein 53 (*Trp53*) (B) cyclin dependent kinase inhibitor 1A (*Cdkn1a*) (C) hypoxia inducible factor 1, alpha subunit (*Hif1a*) (D) NADPH oxidase (*Nox2*) (E) chemokine (C-C Motif) ligand 2 (*Ccl2*) (F) chemokine (C-X-C Motif) ligand 1 (*Cxcl1*) (G) interleukin 6 (*Il6*) (H) interleukin 1 beta (*Il1b*) mRNA levels. Plasma of *Mdm2*cKO (n=4) vs control mice (n=4) was analyzed at day 6 of Dox administration: (I) neutrophils to Lymphocytes ratio (NLR) (CBC Analysis, VetScan Analyzer) (J) IL6 protein levels in plasma. Graphs display mean values ± SEM. Two-tailed t-tests with significance levels *p < 0.05, **p < 0.01, and ***p < 0.001.

As kidney inflammation can contribute to systemic inflammation through the release of inflammatory mediators (18), we found an elevated neutrophil-to-lymphocyte ratio in the *Mdm2*cKO mice compared to the control group (p=0.017) **(Figure 2I)** and a significant increase in plasma IL6 protein levels **(Figure 2J)**. Together, our data indicate systemic inflammation in mice due to severe kidney tubular cell death and kidney dysfunction.

Bulk metabolomics in the kidney cortex of *Mdm2*cKO vs. control mice, revealed prominence of Trp metabolism towards the KP, with significant increases in KYN, 3HK, and QA concentrations (**Figure 3, A-E).** Matrix-assisted laser desorption/ionization-mass spectrometry imaging (MALDI-MSI) confirmed the accumulation of KYN and QA in the kidney, as shown in (**Figure 3, F**-**I**). Upon overlaying ion to autofluorescence images, we observed a diffuse QA expression in both glomerular and non-glomerular regions across the MDM2cKO kidney sections.

**Figure 3:**
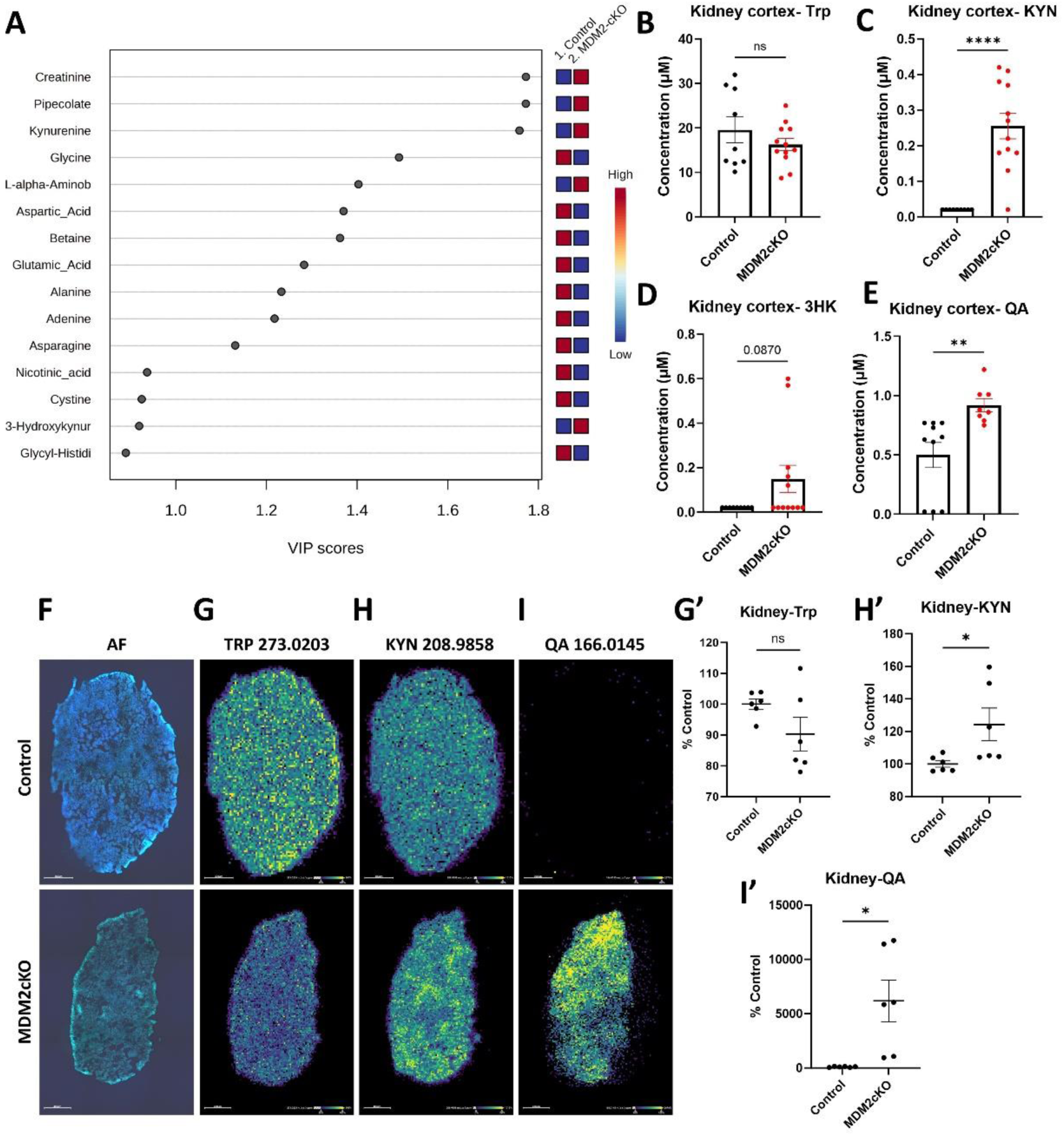
Tryptophan metabolism is skewed in the kidney cortex towards accelerated kynurenine degradation. Targeted metabolomics (CE-MS ZipChip), in kidney cortex tissue from *Mdm2*cKO mice (n=12) compared to control mice (n=9) at day six of Dox administration. (A) Variable importance in projection (VIP) scores from partial least squares discriminant analysis (PLS-DA) of kidney metabolomics. Raw concentrations of (B) tryptophan (Trp); (C) kynurenine (KYN); (D) 3-hydroxykynurenine (3HK); (E) quinolinic acid (QA). (F) Autofluorescence image of control (top) vs. MDM2cKO kidney section (bottom). MALDI-MSI of kidney sections performed in duplicates from *Mdm2*cKO (n=3) vs. control (n=3) mice showing (G) Trp (m/z 273.0203), (H) KYN (m/z 208.9858), and (I) QA (m/z 166.0145), (G’-I’) with their respective semi-quantifications displayed as %control fold change. Pixel intensity blue (low) to yellow (high). Graphs display means ± SEM with significance levels set at *p < 0.05, **p < 0.01, and ****p < 0.0001 via two-tailed t-test.

### Renal dysfunction is associated with increased inflammatory profile and cell death in cortical brain regions

The majority of KYN within the brain comes from circulation via transport across the blood brain barrier (19, 20). As inflammation triggers metabolism of kynurenine within the brain to generate oxidative and neurochemically active metabolites (20–22), we performed targeted bulk metabolomics analysis in the brain cortex of *Mdm2*cKO vs. control mice, focusing on tryptophan metabolites. There were elevated levels of Trp, KYN, 3HK and QA in the brain cortex and a decrease in gamma-aminobutyric acid (GABA) (**Figure 4, A-E)**. Pathway analysis further highlighted Trp and glutamate metabolism in addition to ammonia recycling pathways suggesting a mechanistic link to neuroinflammatory processes (**Supplemental Figure 4 and Supplemental Table 2)**. Perturbed Trp metabolism was accompanied by significant changes in transcripts suggestive of a shift towards cell cycle arrest and inflammation as evidenced by increased *Cdkn1a, Cxcl1, Il1b* mRNA level in the brain cortex of *Mdm2*cKO mice (**Figure 4, F-J)**.

**Figure 4:**
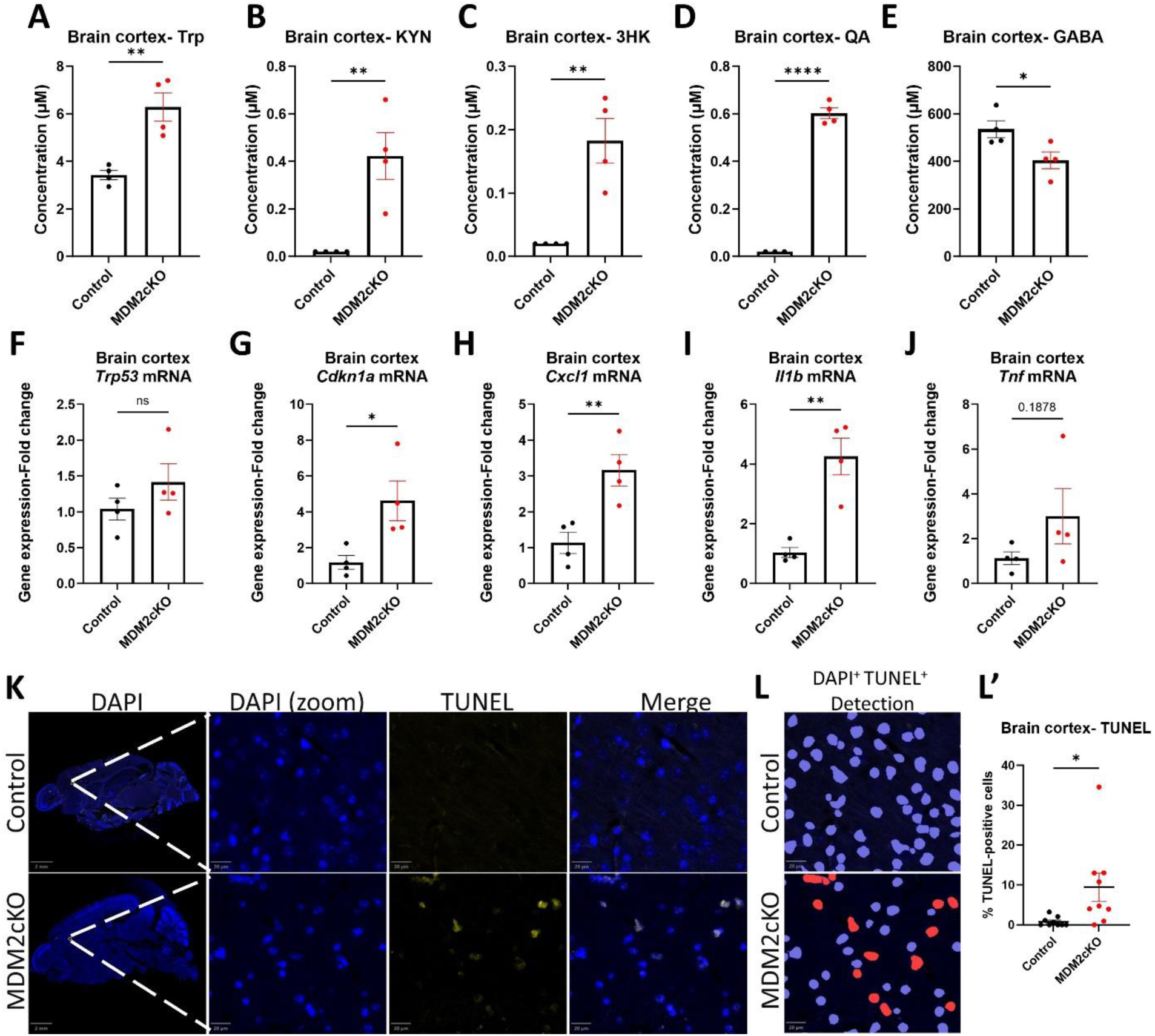
Brains of mice with renal tubular cell death exhibit accelerated kynurenine degradation. Targeted CE-MS ZipChip Bulk metabolomics analysis in brain cortex tissue from *Mdm2*cKO (n=4) vs. control (n=4) mice. Raw concentration displayed in mean+/- SEM for (A) tryptophan (Trp); (B) kynurenine (KYN); (C) 3-hydroxykynurenine (3HK); (D) quinolinic acid (QA); (E) Gamma-aminobutyric acid (GABA). RNA lysates were collected from the brain cortex of *Mdm2*cKO (n=4) vs. control (n=4) mice. qPCR analysis (normalized to GAPDH) showing mRNA fold change in (F) *Trp53*, (G) *Cdkn1a*, (H) *Cxcl1*, (I) *Il1b*, and (J) tumor necrosis factor (*Tnf*). Tunnel assay was conducted on brain sagittal formalin-fixed paraffin-embedded (FFPE) sections obtained from *Mdm2*cKO (n=3) vs. control (n=3) mice. (K) The panels on the left display sagittal brain sections of control (top) vs. *Mdm2*cKO (bottom) mice, white squares indicate a region of interest magnified as indicated by the dashed lines, showing DAPI (λex = 359 nm) in blue and Tunnel (λex = 594 nm) positive cells in yellow, and merged magnified images. (L) QuPath cell detection imaging of nuclei (blue) and TUNEL-positive cells in red. (L’) Graphical representation of % Tunnel-positive cells in the *Mdm2*cKO vs. control groups. Each data point represents an ROI in the brain cortex, 3 ROIs per sample. Two-tailed t-tests, with significance levels *p < 0.05, **p < 0.01, and ****p < 0.0001.two-tailed t-test.

### Spatial metabolomics identified qunilinic acid to be localized in ependymal cells with kidney failure

MALDI-MSI analysis was performed on sagittal fresh frozen brain sections in control and MDM2cKO mice with kidney failure. Gamma-aminobutyric acid (GABA) was expressed in the caudate/striatum and thalamus of the control mice brains and was decreased within the *Mdm2*cKO mice brain sections (**Figure 5 E)**. In contrast, there was a significant increase in KYN levels specifically across grey matter brain regions (**Figure 5 C/C’**). QA was not detectable in control brains by MALDI-MSI, however the MDM2cKO mice brains exhibited significant increased QA intensity in the brain sections of all three analyzed brain samples. Bulk metabolomics confirmed the MALDI-MSI data in (**Figure 4 D).** Further analysis in one of the *Mdm2*cKO mouse brains demonstrated specific localization of QA adjacent or within ependymal cells, which line the cortical ventricle. This was verified by ion image overlay with optical images to clearly demonstrate colocalization of QA with the ependymal cells in post-MALDI H&E-stained tissue (**Figure 5 D/D’ and E-G).**

**Figure 5:**
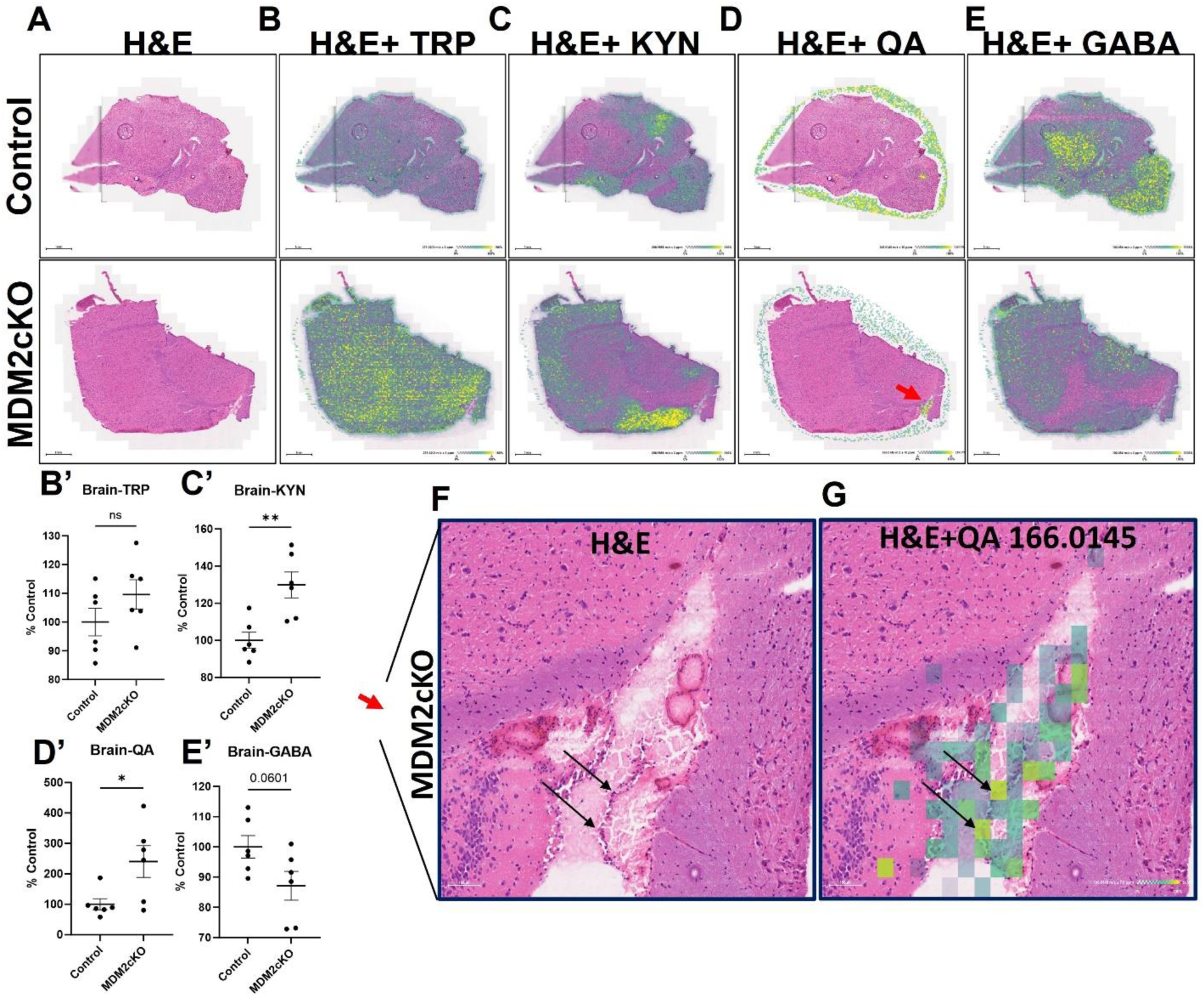
Tryptophan metabolites annotations in MDM2cKO vs. control mice brains exhibit increased kynurenine and quinolinic acid expression. MALDI-MSI were performed in duplicates on brain sagittal sections from n=3 control and n=3 MDM2cko mice. (A) H&E staining overlay with ion image of (B) tryptophan (TRP) (m/z: 273.0203), (C) kynurenine (KYN) (m/z: 208.9858), (D) quinolinic acid (QA) (m/z: 166.0145), and (E) gamma-aminobutyric acid (GABA) (m/z: 102.05605). The red arrow (D) is zoomed in to highlight MDM2cKO mouse brain region of interest (F) H&E, and (G) overlay of H&E with QA ion image; black arrows indicate localization with ependymal cells. (B’-E’) representing respective intensities semi-quantifications of (B-E respectively) displayed in %control change. Graphs display means ± SEM with two-tailed t-tests with *p < 0.05, **p < 0.01.

These metabolic alterations were accompanied by observable changes in cell death at the brain cortical area, underscoring the potential impact of 3HK and QA on brain health. The percentage of TUNEL-DAPI positive cells/ total DAPI-positive cells was significantly increased in the *Mdm2*cKO mice brain cortex compared to control **(Figure 4, J and L/L’)**.

### Accelerated oxidative kynurenine degradation pathway is observed in the ischemia reperfusion (IR) model of severe kidney injury

In order to confirm that the changes in the brain inflammatory signaling milieu that we observed in the *Mdm2*cKO mice could be due to common types of severe kidney tubular injury, we analyzed plasma samples of a mouse model of severe ischemic kidney injury. With 35 minutes of ischemia followed by 24 hours reperfusion (IR35), mice exhibited significant increase in BUN and plasma creatinine level. Interestingly, the IR35 mice plasma exhibited a decrease in Trp and 5-HT and an increase in KYN and 3HK, similar to Trp patterns in plasma of *Mdm2*cKO mice **(Figure 6, A-D).** Moreover, the KYN/Trp ratio strongly correlated to plasma creatinine levels **(Figure 6, E-F)**. Brain analyses in the IR35 mice of cortical regions showed increase in mRNA levels of *Il1b*, *Ccl2* and *Tnf* **(Figure 6, G-I).**

**Figure 6:**
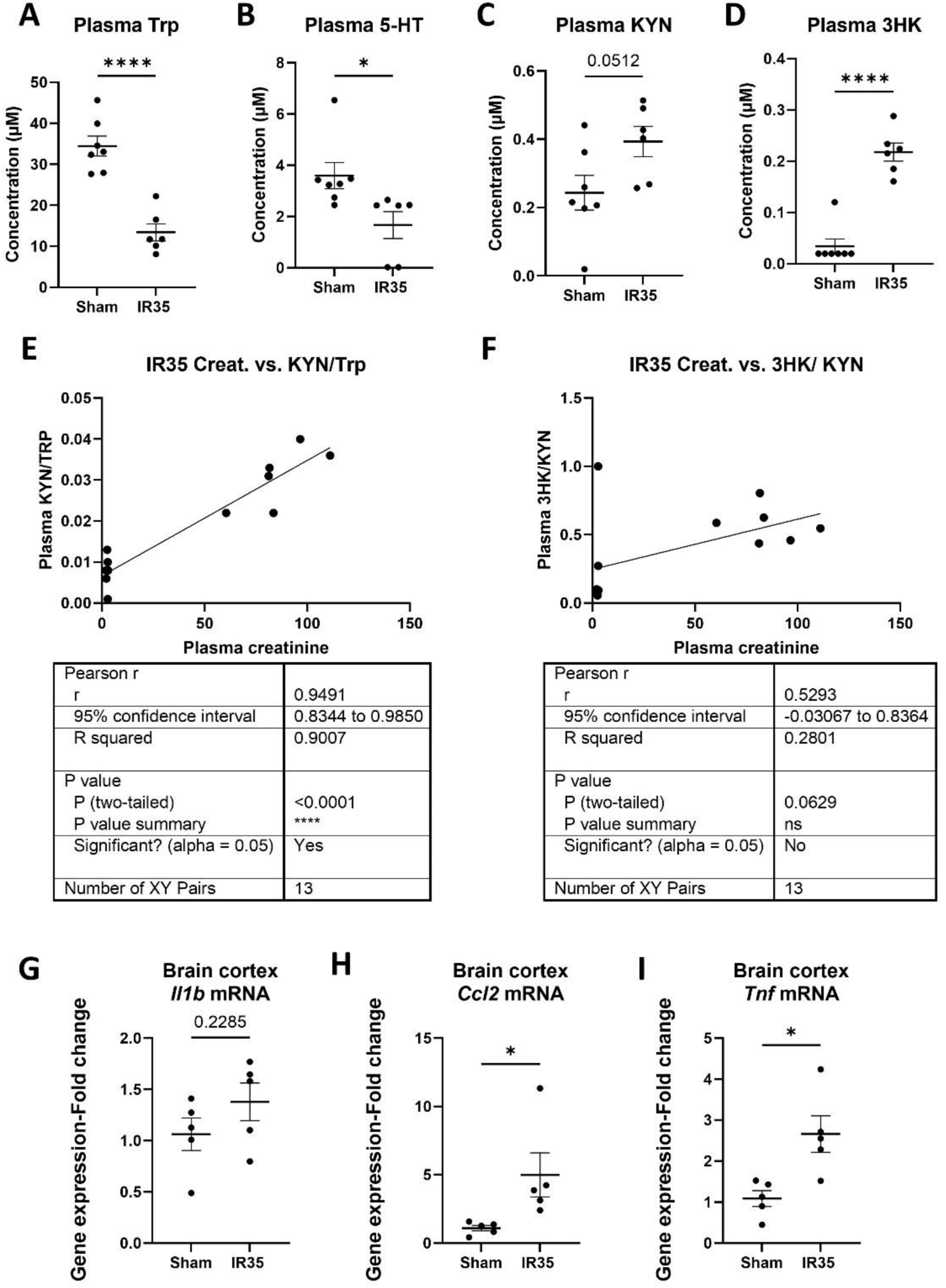
Increased inflammation and accelerated tryptophan degradation is also observed in mouse model of severe kidney injury. Mice were subjected to kidney vasculature clamping for 35 minutes (IR35) sacrificed 24 hours post-reperfusion, and CE-MS ZipChip measurements were performed on plasma samples of IR35 mice vs control mice (Sham). (A-D) Raw concentration of tryptophan (Trp), serotonin (5-HT), kynurenine (KYN), and 3-hydroxykynurenine (3HK). (E-F) Pearson correlations between KYN-to-Trp and 3HK-to-KYN ratios with plasma creatinine. qPCR results for brain cortex RNA lysates, depicting the expression levels of (G-I) *Il1b*, *Ccl2*, and *Tnf*. Graphs display means ± SEM and two-tailed t-test (*p < 0.05, and ****p < 0.0001).

### Advanced kidney disease in humans is associated with increased tryptophan degradation via the kynurenine pathway

In a cohort of CKD patients with a primary etiology of type 2 diabetes and/or hypertension “**Supplemental tables 5-8**”, we observed that levels of plasma Trp were decreased and levels of QA and 3HK were elevated in patients with CKD stage 5 not on dialysis (p <0.01 and p=0.057, respectively), compared with CKD stages 3B and 4 (**Figure 7, A-C)**. These findings, based on a sample size of eight patients with CKD stage 5 and eighteen with stages 3B and 4, indicated a potential rise in neurotoxins associated with the progression of kidney disease severity, similar to the mouse model. Plasma QA was inversely correlated with the estimated glomerular filtration rate (eGFR) and directly correlated with serum creatinine levels in CKD patients (**Figure 7, D and E)**. This suggests that as kidney function declines in humans, there is a concurrent accumulation of QA.

**Figure 7.**
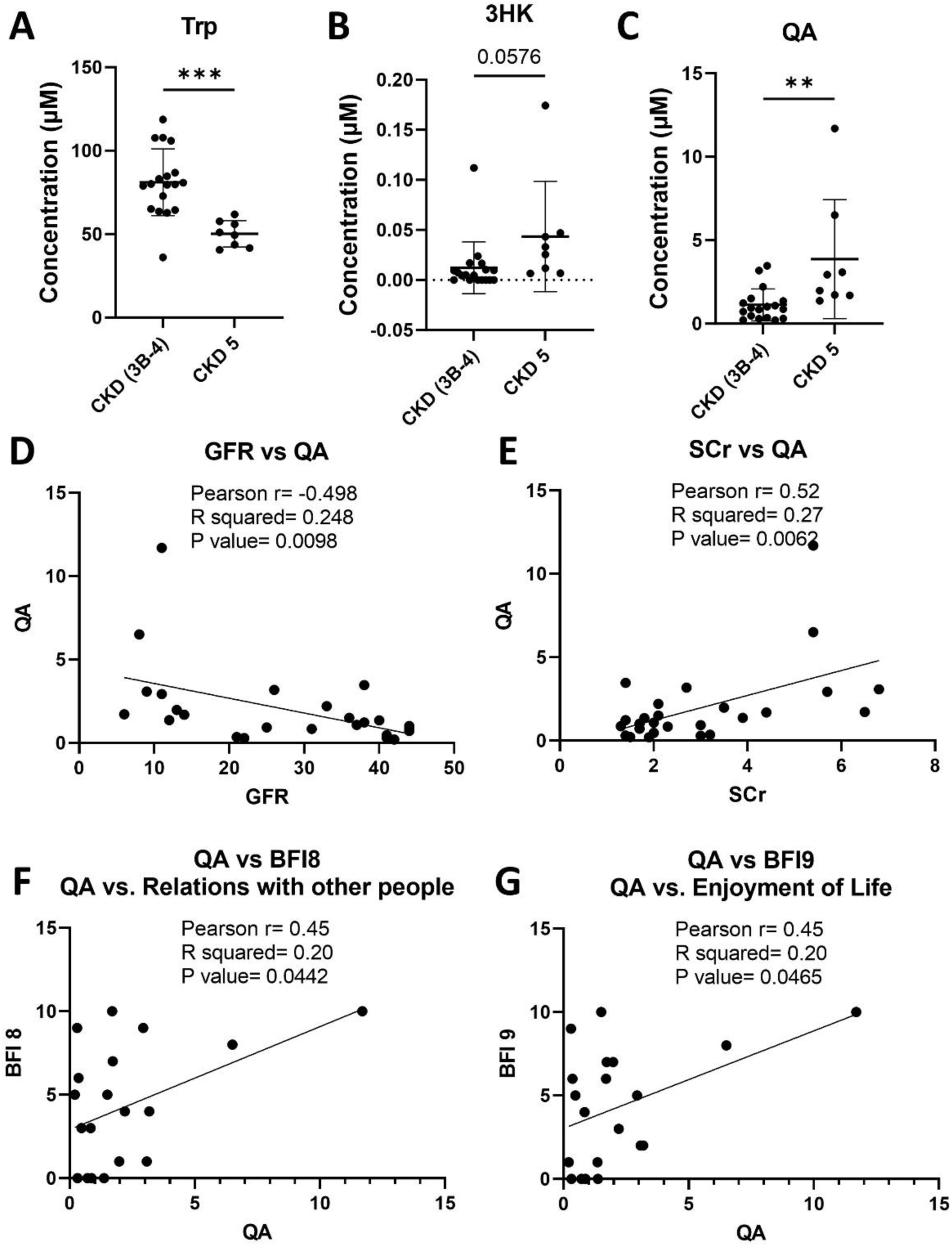
Advanced kidney disease is associated with increased tryptophan degradation via the kynurenine pathway and quinolinic acid inversely correlates with eGFR in chronic kidney disease patients (not on dialysis). Plasma tryptophan metabolites were quantified using LC-MS, n=8 for CKD stage 5 and n=18 for CKD stage 3b and 4. (A) tryptophan (Trp); (B) 3-Hydroxykynurenine (3HK); (C) Quinolinic Acid (QA); (D) Pearson correlation of QA vs. GFR; (E) Pearson correlation of QA vs serum creatinine; (F) Pearson correlation of QA vs interference of fatigue in relations with other people; (G) Pearson correlation of QA vs interference of fatigue in enjoyment of life. Graphs display mean ± SD with statistical significance set via two-tailed t-test with p-values ** p < 0.01 *** p < 0.001.

Examination of the relationship of psychosocial measures and Trp metabolites revealed that higher QA levels were associated with relationships with others (r=0.45, p=0.0442) and enjoyment of life (r=0.45, p=0.0465) (**Figure 7, F and G)**. Correlation analysis did not reveal association with other Trp metabolites, and we could not find any association with other brain fatigue inventory (BFI) items (data not shown), moreover individual BFI item score did not differ between patients with CKD states 3B-4 and stage 5. These findings suggest that the metabolic disturbance in CKD is particularly associated with QA and that QA may have a profound impact on the quality of life (QoL).

## Discussion

In two mouse models of severe kidney dysfunction and in patients with advanced CKD, we observed similar accumulation of downstream bioactive Trp metabolites in the blood. We report that *Mdm2*cKO mice exhibited kidney and systematic inflammation consequent to massive kidney tubular cell death. The metabolic shift in Trp degradation toward the oxidative kynurenine pathway is likely driven by the significant inflammatory response of the damaged kidney at the local and systemic level. It is noteworthy that targeting the kynurenine pathway has been shown to improve kidney injury in KMO null mice (23). Inflammation leading to Trp oxidative degradation is widely supported by previous studies (24–26). This shift has profound implications as it can lead to the production of various neurotoxic and immunoregulatory metabolites. It is notable, that the accumulation of 3HK and QA correlates with risk of developing cognitive impairment and dementia (27, 28), and several neurological disorders wherein altered inflammatory signaling is a key component (22, 26, 29, 30).

Combining data from the *Mdm2*cKO mice model of tubular cell death and the ischemia reperfusion model validate an causative role for tubular injury leading to systemic inflammation and kynurenine degradation. Previous studies have demonstrated a dualistic nature of tubular cells, which can be both susceptible to systemic inflammation and actively contribute to the generation of inflammatory mediators (18). These processes can significantly impact the progression of kidney injury and its impact on distant organs.

### Tryptophan metabolism via the kynurenine pathway in the brain is associated with increase in QA, inflammatory signaling changes and increased cell death

Our pathway enrichment analysis from the targeted bulk metabolomics performed on the brain cortical lysates of *Mdm2*cKO mice highlighted tryptophan and glutamate pathways among the top 5 significantly skewed pathways. The interaction between tryptophan metabolism through the oxidative kynurenine pathway and glutamate metabolism in the brain presents an intriguing avenue for exploration. Oxidative kynurenine pathway is driven by inflammation in the brain (31) our data in both mouse models and prior research document altered brain inflammatory profile under kidney injury (32). Interestingly, QA is a bioactive kynurenine oxidative byproduct, typically found in very low nanomolar levels in the human brain and cerebrospinal fluid (CSF), however QA can reach toxic levels in response to inflammation. Similarly our quantitative bulk metabolomic analysis demonstrated undetectable QA in the control brains vs an average of 0.6 µM QA in the MDM2cKO brain cortex. QA is frequently referred to as an excitotoxic or neurotoxic brain metabolite and has been associated with the development of various neurological diseases in humans (21). QA interacts with NMDA receptors, competing with glutamate and contributes to excitotoxicity and neuronal damage (21, 33–36). Moreover, our finding of decreased GABA levels in the brain cortex is consistent with other studies showing GABA as a suppressor of inflammation and excitotoxic damage (37, 38) and its decrease in the brain is associated with various neurological disorders (39, 40)

Our novel spatial metabolomics approach overlaying MALDI-MSI ion images to post-MALDI H&E-stained brain tissue evealed an interesting pattern wherein QA colocalized with ependymal cells in one of the MDM2cKO brains. Ependymal cells are glial cells that line the cerebral ventricles and secrete and circulate CSF. Interestingly, CSF QA levels has been significantly correlated with neurological disorders associated with conditions such as HIV infection, traumatic brain disorder and hepatic encephalopathy (41–46). Moreover, the QA catabolic enzyme quinolinic acid phosphoribosyltransferase (QPRTase) was observed via immuno-histochemistry in ependymal cells of the cerebral ventricles (47). Cumulatively our data and published studies indicate that QA is closely implicated with inflammation in the brain and excitotoxicity, and that ependymal cells may play a role in the metabolism or regulation of quinolinic acid in the brain. However, the functional significance of the localization of QA to ependymal cells has not yet been investigated.

### Inflammation, favoring oxidative Trp catabolism could lead to fatigue

A significant percentage of CKD patients not on kidney replacement therapy report fatigue (48, 49). In general CKD patients have a worse quality of life (48, 50). However, the underlying mechanisms are poorly understood. We and others have previously demonstrated a significant inverse association between circulating Trp levels and stages of CKD (24, 51). KYN, kynurenic acid, and QA, were positively and robustly correlated with the severity of kidney disease due to higher enzymatic activity induced mainly by inflammation.

In the present study, we consistently found that patients with CKD stage 5, compared to stages 3b/4, had significantly low circulation plasma free Trp levels coupled with high levels of KP pathway metabolites indicating enhanced IDO activity. Here, we also observed a strong correlation specifically between plasma QA and scores of fatigue interference with certain daily activities, notably relationship with people and enjoyment of life. To our knowledge this is the first data linking QA specifically to psychosocial changes in CKD patients.

Our data are consistent with several reports that showed correlation of QA with fatigue in patients with chronic fatigue syndrome, fibromyalgia, and systemic lupus erythematosus (52, 53). Reducing KYN in postmenopausal breast cancer survivors has demonstrated to correlate with reduced fatigue (54). Our research holds particular relevance for neurologic dysfunction associated with progressive CKD and acute kidney injury (55). Ongoing research into the kidney-brain axis via metabolic alterations arising from the kidney offers exciting prospects for targeted treatments.

## METHODS

### *Mdm2* conditional knock-out mice

*Pax8*-rtTAcre;*Mdm2*f/f or *Mdm2*f/f control 3-5 months old mice, maintained on C57BL/6J mice background, were treated with doxycycline (DOX) 2mg/ml in 5% sucrose drinking water (6-20 days) to induce deletion of *Mdm2* in paired box 8 (*Pax8*)-positive cells (15, 16, 56). Mice were monitored daily during doxycycline administration. Blood was collected via submandibular vein for metabolomics analysis (CE-MS; ZipChip) or complete blood count (CBC) blood analysis - Vetscan-HM5-Hematology Analyzer. Primers for mice genotyping detailed in (**Supplemental Table 4).** *Pax8*-rtTAcre mice were provided as gift from Karen Block, PhD University of Texas Health Science Center at San Antonio and the *MDM2f/f* mice were obtained from MD Anderson (57).

### Ischemia-reperfusion injury (IRI) in mice

Ten-to twelve-week-old male C57BL/6J mice were anesthetized by isoflurane inhalation (3% in oxygen) and body temperature was maintained at 36-37 °C. Bilateral renal ischemia-reperfusion (IR) was conducted as previously described (58) with some modifications. Briefly, the renal pedicles were clamped for 35 minutes with small nontraumatic vascular aneurysm clips (Roboz Surgical Instruments, Gaithersburg, MD). Subcutaneous saline injection of 0.5 ml was administered after performing two-layered incisions. Same surgical procedures, except for pedicle clamping, were applied on Sham-operated mice.

### Targeted bulk metabolomics

Plasma, kidney cortex or brain cortex lysates were analyzed using ZipChip (908 Devices, Boston, MA) coupled with mass spectrometry (17, 59, 60). A microfluidic chip that integrates capillary electrophoresis (CE) with nano-electrospray ionization through a ZipChip interface separates metabolites. Q-Exactive mass spectrometer (Thermo, San Jose, CA) was used for data acquisition, and data processing was done via Thermo Scientific’s software Xcalibur-Quan Browser. A detailed protocol is available (61). Data and pathway analysis were performed using MetaboAnalyst 5.0 (62).

### Matrix-assisted laser desorption ionization mass spectrometry imaging (MALDI-MSI)

Frozen brain or kidney tissue sections from mice were subjected to MALDI-MSI in negative ion mode using an Orbitrap mass spectrometer operated at a resolution of 120,000 at m/z 200. Cryosectioned tissue samples were coated with a matrix compound, and mass spectra were acquired in imaging mode to generate high-resolution ion images, details described in (63). The annotations were processed using CoreMetabolome - v3 and HMDBI on METASPACE. Images were extracted on SCiLS Lab software followed by Total Ion Current (TIC) normalization to account for variations in ion intensity across the tissue sections.

### RNA extraction and quantitative polymerase chain reaction (qPCR)

Total RNA was extracted using QIAGEN RNeasy Mini Kit (Cat# 74104). Briefly, 500 µL lysate was mixed with 500 µL of 70% ethanol and applied to RNeasy Mini spin column. After washing steps, RNA was eluted in 30-50 µL of RNase-free water. The RNA concentration and purity were determined using NanoDrop spectrophotometer. cDNA synthesis was performed using Thermo Fisher Scientific RevertAid Reverse Transcription Kit (Cat# 4374966) with total of 1 µg of RNA for each sample reaction. qPCR master mix was prepared by mixing 5 µL SYBR Green PCR Master Mix (Thermo Fisher Scientific, Cat# A25780) with 0.5 µL primer mix (forward and reverse 10uM) and nuclease-free water to reach a final reaction volume of 11 µL per reaction. The primers sequences are listed in (**Supplemental table 3).**

### Terminal deoxynucleotidyl transferase dUTP Nick-End Labeling (TUNEL) assay

TUNEL assay was performed using the Elabscience (E-CK-A322) kit. Brain tissue samples were collected and stored in 70% ethanol at 4^0^C after 24 hour fixation in 10% formalin, followed by paraffin embedding and sagittal sectioning. Positive and negative control measures were employed. The TUNEL assay results were analyzed using QuPath software (64). Three regions of interests were selected in the cortex of every sample at similar locations from control (n=3) vs. *Mdm2*cKO (n=3). The fluorescent signal was captured using Zeiss Axioscan 7 microscope equipped with a 10x/0.45 Plan Apochromat objective, with excitation wavelengths λex = 359 nm (DAPI) and λex = 594 nm (TUNEL).

### Study population

In this observational study, we enrolled adult CKD patients treated at an adult outpatient nephrology clinic with following eligibility criteria: (i) clinical diagnosis of CKD stages 3B, 4, and 5 (ii) primary etiology of CKD is either type 2 diabetes and or hypertension, and (iii) on standard management for diabetes, hypertension, and associated comorbidities including anemia as per the recommended guidelines(65). The study was approved by the local Institutional Review Board and all participants provided written informed consent prior to the study procedures.

### Data collection

During a routine clinic visit, each participant completed the Brief Fatigue Inventory (BFI) to report fatigue. Random blood was collected from each consented participant in serum separator tube for serum and anticoagulant containing vacutainer for plasma. Each participant also provided random spot urine which was measured for albumin and creatinine as per standard methodologies. Relevant sociodemographic (age, sex, height, and weight), clinical (medical history and concomitant medications) and laboratory data (e.g., glycated hemoglobin, blood hemoglobin) were reviewed and abstracted from the electronic health record. Serum creatinine was measured using a kinetic rate Jaffé method. The estimated glomerular filtration rate (eGFR) was calculated using serum creatinine, age and sex based on the Chronic Kidney Disease Epidemiology Collaboration (CKD-EPI) equation (66). **Supplemental Table 4** shows the stages of chronic kidney disease based on eGFR categories.

### Tryptophan metabolites

Free tryptophan and selective metabolites in the KP were measured in plasma by liquid chromatography/mass spectrometry (LC-MS) as reported previously(31). Briefly, 50μL plasma was diluted with 5 times of 0.2% acetic acid. Stable isotope–labeled standards, 2-picolinic-d4 acid, 2,3-pyridinedicarboxylic acid-d3, L-Trp-13C11,15N2, and KYN, were added at the time of extraction as internal standards for absolute quantification. The diluted samples were vortexed and transferred to 0.5-mL Millipore Amicon Ultra filter (3 kDa). The filter tubes were centrifuged at 13 500g for 60 minutes at 4°C and the extracts were transferred to glass vials for LC-MS analyses. High-performance liquid chromatography/electrospray ionization mass spectrometry (HPLC-ESI-MS) analyses were conducted on a Thermo Fisher Q Exactive mass spectrometer with online separation by a Thermo Fisher/Dionex Ultimate 3000 HPLC. High-performance liquid chromatography conditions were as follows: column, YMC-Pack ODS-AQ, 3 μm, 2 × 100 mm (YMC; Allentown, PA); mobile phase A, 0.5% formic acid in water; mobile phase B, 1% formic acid in acetonitrile; flow rate, 200 μL/min; gradient, 1% B to 30% B for 5 minutes and held at 70% B for 5 minutes to clean the column. The MS analyses were conducted using full MS scan (70 000 resolution) with positive ion detection. Standard curves were generated for all targeted KYN compounds using appropriate stable isotope–labeled internal standards and native compounds. Quantitative results were obtained by reference of the experimental peak area ratios to the standard curves. **Brief Fatigue Inventory (BFI):** BFI is a 9-item questionnaire that measures fatigue during the past 24-hour on a 0-10 Likert scale with higher scores representing worse fatigue (67). The first three BFI items measure fatigue severity with score ranging from 0 (no fatigue) to 10 (fatigue as bad as you can imagine). The remaining six BFI items assess fatigue interference in relation to patients’ general activity, mood, walking ability, normal work (both indoor and outdoor), relations with other people, and enjoyment of life. These fatigue interference items represent the pervasive impact of fatigue on daily life activities – one of the most important and prioritized outcomes in patients with CKD (68). Fatigue interference items are measured on a 0–10 numerical rating scale, with 0 being “does not interfere” and 10 being “completely interferes.” **Blinded research:** Metabolomics measurements (bulk and spatial) were conducted blindly by research technician experts without access to samples’ groups identification.

### Sex as a biological variable

In the human and most of the mouse studies, sex was not considered as biological variable. In the IRI studies only males were employed because females have significantly increased tolerance to IRI (69).

### Statistics

For human studies, descriptive data are presented as mean +/- SD, and all comparisons are two-tailed unpaired t-test. We performed Pearson correlation coefficient test to measure the association between two variables. For mouse studies two-tailed unpaired t-test for comparison between groups, were employed and data presented as mean +/-SEM. Moreover, Pearson correlation test was performed. All analyses were performed on GraphPad Prism 8 with p-values < 0.05 reported as significant. In addition, MetaboAnalyst 5.0 was used for metabolic data analysis, principal component analysis (PCA), and partial least squares discriminant analysis (PLS-DA) to scrutinize metabolites differences between groups. For metabolic enrichment analyses SMPDB (The Small Molecule Pathway Database) weas employed to recognize metabolic pathways significance reported with adjusted FDR p-values <0.05.

### Study approval

For human studies, all study procedures including biospecimen collections were performed after obtaining written consent of IRB approved study protocol from each participant (IRB approval 20140210HU). Mouse studies were carried out after IACUC approval from the University of Texas Health Science Center at San Antonio.

### Data availability

All the data necessary to support the conclusions presented in this manuscript are included within the figures, tables, or supplemental material unless clearly stated otherwise.

## Author contributions

AS and KS conceptualized the research. AS designed and conducted the animal research, acquired and analyzed data. SD designed and performed human research. IT performed the MALDI-MSI analysis. JT and MM contributed to the ischemia reperfusion studies in mice. PSM contributed to mice samples acquisition. CF contributed to brain histology. SM and HJL contributed with quality standard operating procedures; GZ contributed with metabolomics pathway analysis; LH ensured quality control of mass spectrometry data analysis. AS received scientific guidance and insights from SD (clinical data, CKD and tryptophan metabolism), JCO (Inflammation, tryptophan metabolism, neuroscience), BF (data analysis, neuroscience), SCH (brain histology, inflammation), KFB (brain histology and interpretation), JDL (neuroscience). AS wrote the manuscript. All authors reviewed, provided edits and approved the submission of the manuscript. KS provided oversight, scientific and grant acquisition support for all the work.

## Acknowledgments

We thank the team of the Center for Precision Medicine at the University of Texas Health Science Center at San Antonio (UTHSA) for peer review, scientific discussions, and technical support, namely, Shane Matta, M.Sc. and Anthony Franzone for acquisition of mass spectrometry data.

## This research was supported by

National Institutes of Health, National Center for Advancing Translational Sciences grant TL1 TR002647 (AS)

National Institutes of Health, National Institute of Diabetes and Digestive and Kidney Diseases UO1, 5U01DK114920-06; Department of Defense W81XWH1910659, VA Merit Review #2I01BX001340-09A1, I101BX003234 (KS)

University of Texas Health Science Center at San Antonio, Long School of Medicine, Multi-PI pilot grant 2023-2024 (KS, BF)

National Institutes of Health, National Institute of Neurological Disorders and Stroke, K01NS126489 (BF). In addition, BF is partially supported by the South Texas Alzheimer’s Disease Research Center (P30AG066546).

National Institutes of Health, National Institute of Aging, K01-AG066747 (SCH) and T32-AG021890 (CF).

The content is solely the responsibility of the authors and does not necessarily represent the official views of the NIH.

