## Supplemental File for "Quinolinic acid links kidney injury to brain toxicity"

**Supplemental Figure 1:** Characterization of Pax8-rtTAcre;MDM2f/f mouse model.

**Supplemental Figure 2:** Comparative CE-MS analysis of metabolites in plasma of Mdm2cKO vs. control mice following doxycycline administration.

**Supplemental Figure 3:** Partial least squares discriminant analysis (PLS-DA) principal component analysis (PCA) of bulk metabolomics in plasma, kidney cortex, and brain cortex of MDM2cKO vs. control mice.

**Supplemental Figure 4:** Metabolic pathway analysis reveals dysregulation in tryptophan and glutamate metabolism in MDM2cKO mouse brain cortex.

**Supplemental Figure 5:** MDM2 protein and mRNA Levels in the brain lysates of MDM2cKO vs. control mice.

**Supplemental Table 1:** Results from quantitative enrichment analysis using MetaboAnalyst 5.0 with MBRole2.0 open databases, incorporating SMPDB - CE-MS targeted bulk metabolomics of 32 metabolites in plasma of MDM2cKO vs. control mice.

**Supplemental Table 2:** Results from quantitative enrichment analysis using MetaboAnalyst 5.0 with MBRole2.0 open databases, incorporating SMPDB - CE-MS targeted bulk metabolomics of 32 metabolites in brain cortex of MDM2cKO vs. control mice.

**Supplemental Table 3:** qPCR primers list

**Supplemental Table 4.** Primers list for mouse genotyping

**Supplemental Table 5:** Stages of chronic kidney disease based on eGFR categories (mL/min/1.73m$^2$) description and range

**Supplemental Table 6:** Characteristics of the study participants

**Supplemental Table 7:** Plasma-free levels of tryptophan and selective tryptophan metabolites and ratios

**Supplemental Table 8:** Self-reported scores of brief fatigue inventory

**Supplemental Methods**

**Supplemental Figure 1: Characterization of Pax8-rtTAcre;MDM2f/f mouse model.** (A) Schematic representation of experimental design in Pax8-rtTAcre;MDM2f/f and control MDM2 fl/fl mice, both administered doxycycline (Dox) (2 mg/ml in 5% sucrose drinking water) for tissue and specimen collection at day 6. (B) Body weight loss at day 6 of doxycycline treatment (n=4 per group). (C) Survival percentage across 15 days of doxycycline n=10 per group (D) Hematoxylin and Eosin (H&E) staining of kidney FFPE sections, comparing control and MDM2cKO mice kidneys at X10 and X40 magnifications using brightfield microscopy. (E) Blood urea Nitrogen (BUN) levels at day 3 and day 6 of Dox administration. (F) Plasma creatinine levels at day 3 vs. day 6 of Dox administration, with n=3 control and n=4 MDM2cKO. (One-way ANOVA test ***p < 0.001 and ****p < 0.0001.) (G) Schematic illustrating the extraction of renal proximal tubule epithelial cells from Pax8-rtTAcre;MDM2f/f or MDM2 fl/fl mice, followed by treatment with Dox at 1 µg/ml, and cell harvest at 24 hours. (H) mRNA levels fold change of MDM2 normalized to GAPDH. Schemes created with Biorender.com.

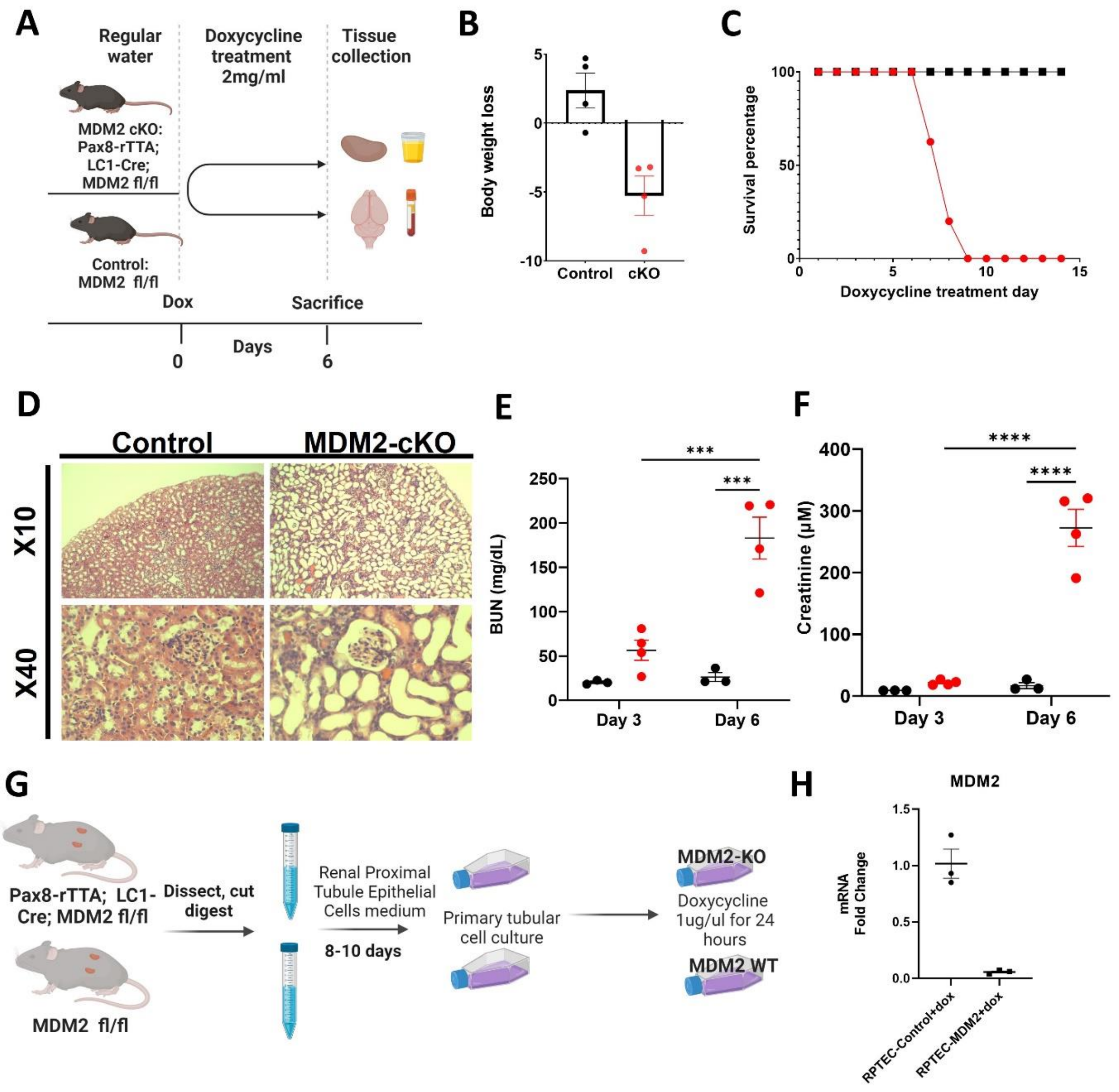

**Supplemental Figure 2: Comparative CE-MS analysis of metabolites in plasma of Mdm2cKO vs. control mice following doxycycline administration at day 3 and day6.** (A) Tryptophan; (B) Kynurenine; (C) 3-Hydroxykynurenine (3HK); (D) Pearson correlation analysis of plasma kynurenine-to-tryptophan (KYN/Trp) ratio vs. plasma creatinine levels across day 3 and day 6. Graph display means ± SEM with one-way ANOVA test with significance levels set at *p < 0.05, **p < 0.01, ***p < 0.001, and ****p < 0.0001. (E) Pearson correlation analysis of plasma 3-hydroxykynurenine-to-kynurenine (3HK/KYN) ratio versus creatinine levels across day 3 and day 6 of Dox administration. Correlation analyses include both MDM2cKO and control groups.

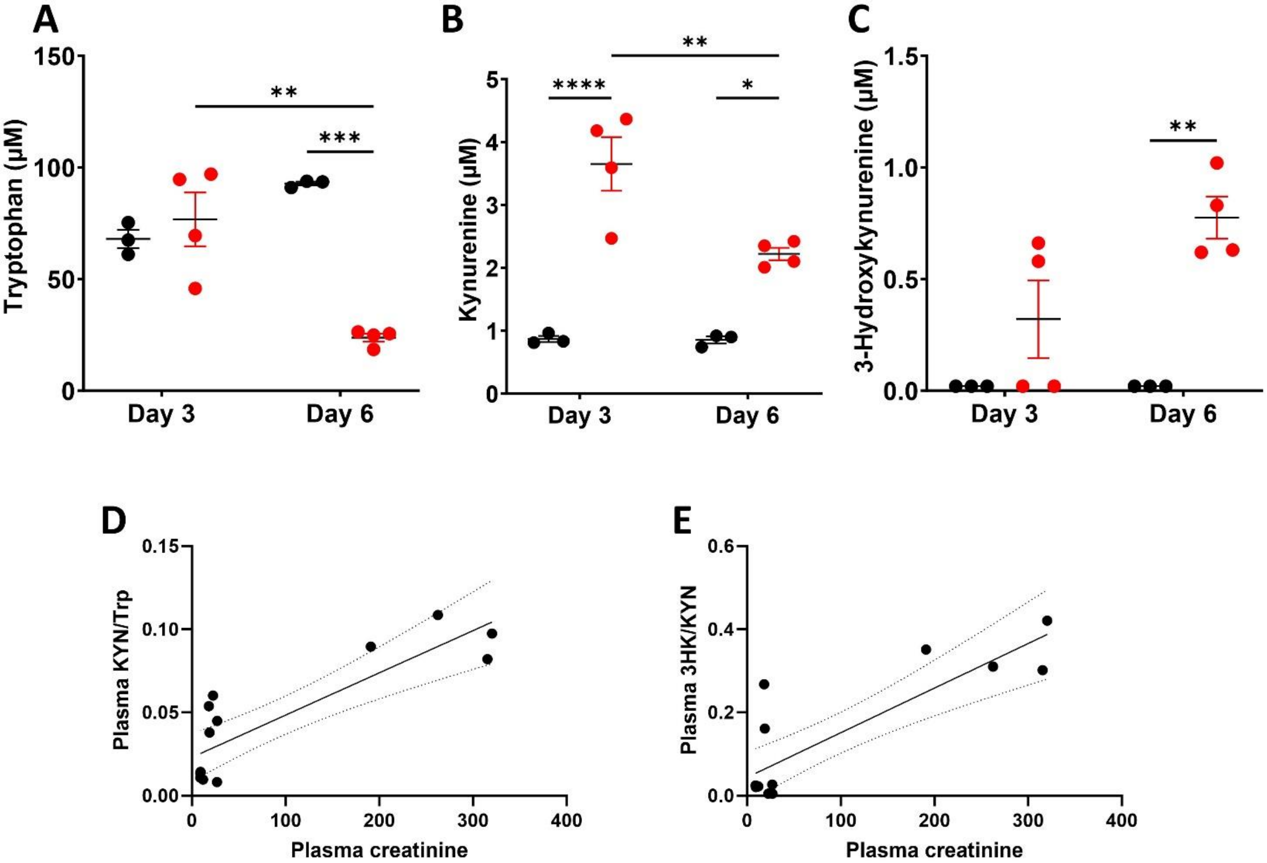

| Pearson r | |
|---|---|
| r | 0.8562 |
| 95% confidence interval | 0.5967 to 0.9536 |
| R squared | 0.7331 |
| P value | |
| P (two-tailed) | <0.0001 |
| P value summary | **** |
| Significant? (alpha = 0.05) | Yes |
| Number of XY Pairs | 14 |

| Pearson r | |
|---|---|
| r | 0.8527 |
| 95% confidence interval | 0.5882 to 0.9524 |
| R squared | 0.7271 |
| P value | |
| P (two-tailed) | 0.0001 |
| P value summary | *** |
| Significant? (alpha = 0.05) | Yes |
| Number of XY Pairs | 14 |

**Supplemental Figure 3: Partial least squares discriminant analysis (PLS-DA) principal component analysis (PCA) of bulk metabolomics in plasma, kidney cortex, and brain cortex of MDM2cKO vs. control mice.** Data from bulk targeted metabolomics of 32 metabolites of MDM2cKO vs. control mice were log-transformed and autoscaled to account for differences in metabolite concentrations. (A) plasma; (B) kidney cortex and (C) brain cortex. PLS-DA PCA plots were generated to show the separation between MDM2cKO mice and control mice based on their metabolic profiles. Each data point represents an individual mouse, with MDM2cKO mice shown in green and control mice shown in red. (A') plasma; (B') kidney cortex; (C') brain cortex.

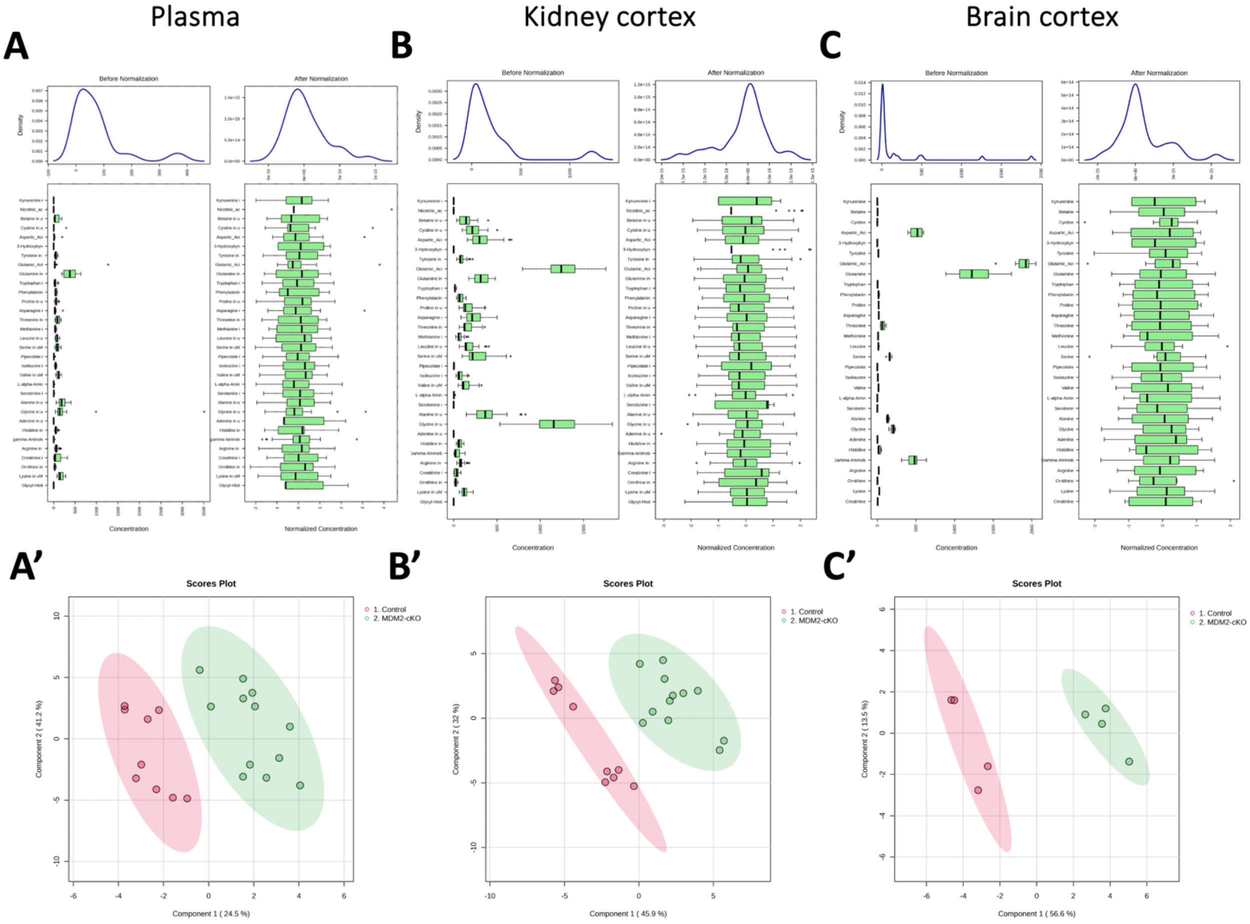

**Supplemental Figure 4: Metabolic pathway analysis reveals dysregulation in tryptophan and glutamate metabolism in MDM2cKO mouse brain cortex.** (A) Targeted metabolomics and pathway enrichment analysis in the brain cortex of MDM2cKO mice vs. control mice involving 32 metabolites. An overview of unbiased pathway enrichment analysis was performed using MetaboAnalyst 5.0 with MBRole2.0 open databases using SMPDB library. (B) Threonine and 2-oxobutanoate Degradation: 1 significant hit, FDR < 0.001. (C) Tryptophan metabolism: 4 significant hits out of 5 were identified, FDR < 0.001. (D) Glutamate metabolism: 3 significant hits out of 6, FDR < 0.01. (E) Ammonia metabolism: 3 significant hits out of 6, FDR < 0.01. (F) Purine metabolism: 2 significant hits out of 5, FDR < 0.01.

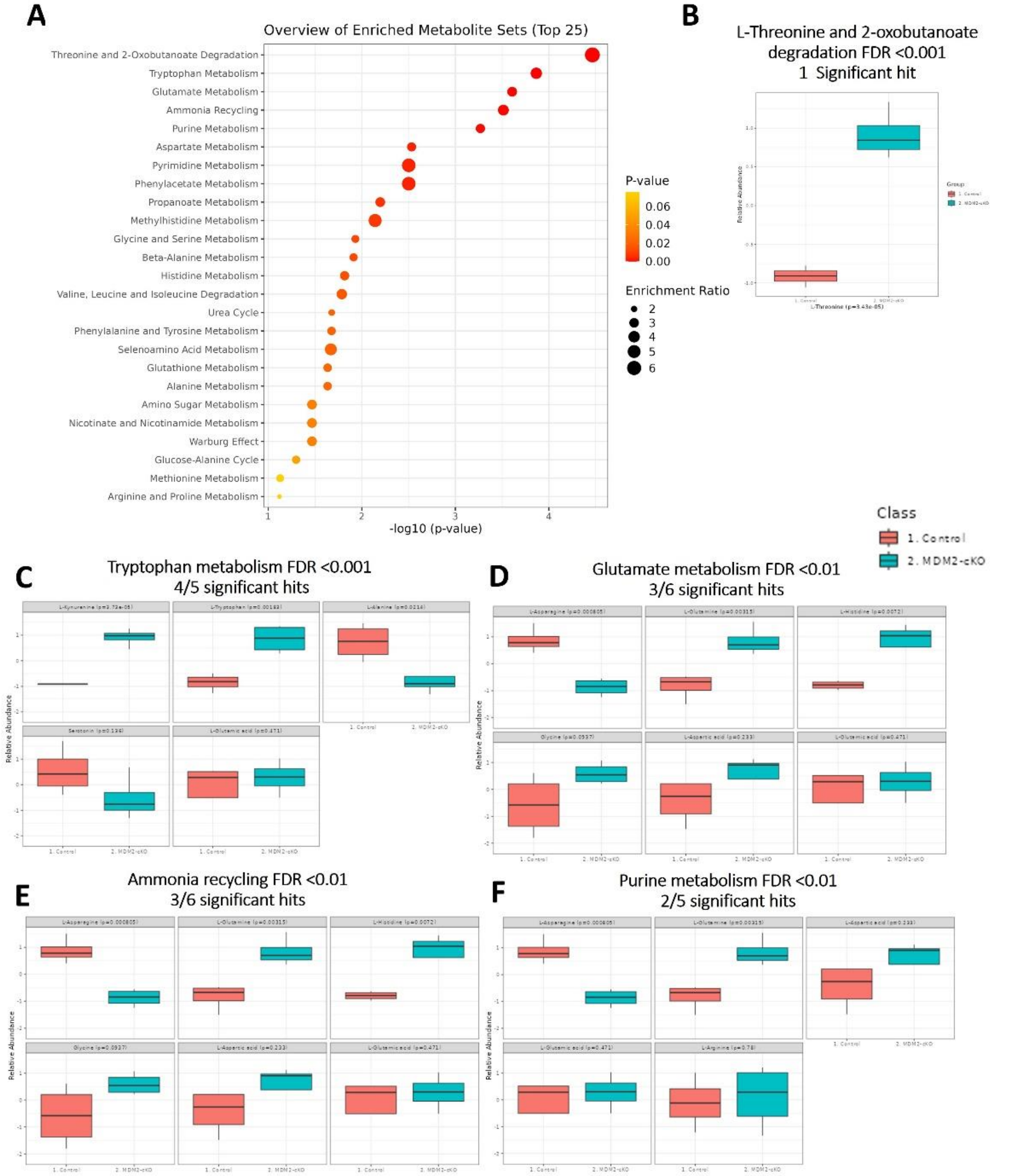

**Supplemental Figure 5: MDM2 protein and mRNA Levels in the brain lysates of MDM2cKO vs. control mice.** (A) Western blot analysis used 10 µg/well of brain lysate extracted with RIPA buffer. LICOR imaging of MDM2 protein (54 KDa) in red using IRDye 680RD (emission at 680 nm) and beta-actin (42 KDa) in green with IRDye 800CW (emission at 792 nm). The full, unedited blot is presented for n=4 MDM2-cKO mice vs. n=4 control mice. (A') Quantitative analysis of Western blot data, with values normalized to beta-actin levels. (B) qPCR analysis of brain cortex *Mdm2* mRNA normalized to GAPDH mRNA. Graph display means +/- SEM and two-tailed t-test, with * $p < 0.05$; *** $p < 0.001$.

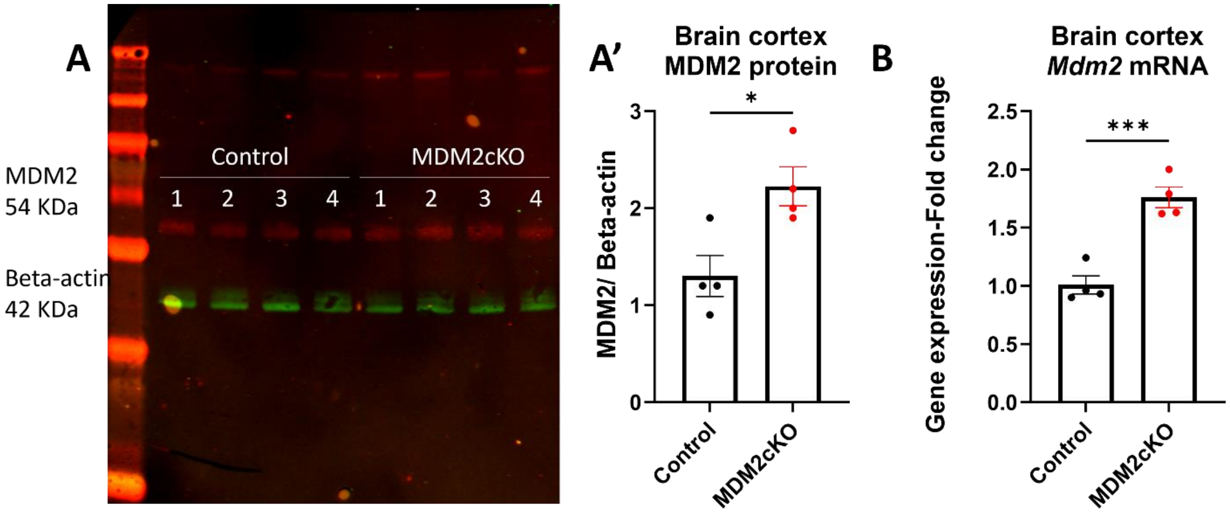

**Supplemental Table 1.** Results from quantitative enrichment analysis using MetaboAnalyst 5.0 with MBRole2.0 open databases, incorporating SMPDB - CE-MS targeted bulk metabolomics of 32 metabolites in plasma of MDM2cKO vs. control mice.

| | Total Cmpd | Hits | Statistic Q | Expected Q | Raw p | Holm p | FDR |
|---|---|---|---|---|---|---|---|
| Tryptophan Metabolism | 60 | 5 | 28.91 | 5.00 | 1.93E-04 | 7.52E-03 | 7.52E-03 |
| Selenoamino Acid Metabolism | 28 | 1 | 25.03 | 5.00 | 2.09E-02 | 7.95E-01 | 2.52E-01 |
| Methylhistidine Metabolism | 4 | 1 | 24.37 | 5.00 | 2.30E-02 | 8.50E-01 | 2.52E-01 |
| Catecholamine Biosynthesis | 20 | 1 | 21.91 | 5.00 | 3.24E-02 | 1.00E+00 | 2.52E-01 |
| Thyroid hormone synthesis | 13 | 1 | 21.91 | 5.00 | 3.24E-02 | 1.00E+00 | 2.52E-01 |
| Glucose-Alanine Cycle | 13 | 2 | 12.54 | 5.00 | 8.06E-02 | 1.00E+00 | 4.57E-01 |
| Histidine Metabolism | 43 | 2 | 12.21 | 5.00 | 8.20E-02 | 1.00E+00 | 4.57E-01 |
| Spermidine and Spermine Biosynthesis | 18 | 2 | 9.60 | 5.00 | 1.59E-01 | 1.00E+00 | 4.60E-01 |
| Urea Cycle | 29 | 6 | 8.45 | 5.00 | 1.63E-01 | 1.00E+00 | 4.60E-01 |
| Pyrimidine Metabolism | 59 | 1 | 9.86 | 5.00 | 1.66E-01 | 1.00E+00 | 4.60E-01 |
| Phenylacetate Metabolism | 9 | 1 | 9.86 | 5.00 | 1.66E-01 | 1.00E+00 | 4.60E-01 |
| Glutathione Metabolism | 21 | 3 | 8.38 | 5.00 | 1.68E-01 | 1.00E+00 | 4.60E-01 |
| Alanine Metabolism | 17 | 3 | 8.38 | 5.00 | 1.68E-01 | 1.00E+00 | 4.60E-01 |
| Glycine and Serine Metabolism | 59 | 8 | 7.84 | 5.00 | 1.86E-01 | 1.00E+00 | 4.60E-01 |
| Beta-Alanine Metabolism | 34 | 3 | 8.15 | 5.00 | 1.94E-01 | 1.00E+00 | 4.60E-01 |
| Phenylalanine and Tyrosine Metabolism | 28 | 3 | 7.66 | 5.00 | 2.01E-01 | 1.00E+00 | 4.60E-01 |
| Betaine Metabolism | 21 | 2 | 7.98 | 5.00 | 2.07E-01 | 1.00E+00 | 4.60E-01 |
| Glutamate Metabolism | 49 | 5 | 7.01 | 5.00 | 2.23E-01 | 1.00E+00 | 4.60E-01 |
| Tyrosine Metabolism | 72 | 3 | 7.33 | 5.00 | 2.24E-01 | 1.00E+00 | 4.60E-01 |
| Threonine and 2-Oxobutanoate Degradation | 20 | 1 | 5.86 | 5.00 | 2.90E-01 | 1.00E+00 | 5.64E-01 |
| Ammonia Recycling | 32 | 6 | 5.81 | 5.00 | 3.09E-01 | 1.00E+00 | 5.64E-01 |
| Propanoate Metabolism | 42 | 2 | 5.65 | 5.00 | 3.31E-01 | 1.00E+00 | 5.64E-01 |
| Valine, Leucine and Isoleucine Degradation | 60 | 4 | 5.25 | 5.00 | 3.33E-01 | 1.00E+00 | 5.64E-01 |
| Methionine Metabolism | 43 | 3 | 5.34 | 5.00 | 3.51E-01 | 1.00E+00 | 5.65E-01 |
| Amino Sugar Metabolism | 33 | 2 | 4.95 | 5.00 | 3.77E-01 | 1.00E+00 | 5.65E-01 |
| Warburg Effect | 58 | 2 | 4.95 | 5.00 | 3.77E-01 | 1.00E+00 | 5.65E-01 |
| Aspartate Metabolism | 35 | 5 | 4.28 | 5.00 | 4.08E-01 | 1.00E+00 | 5.89E-01 |
| Nicotinate and Nicotinamide Metabolism | 37 | 3 | 4.55 | 5.00 | 4.48E-01 | 1.00E+00 | 6.24E-01 |
| Purine Metabolism | 74 | 5 | 3.97 | 5.00 | 5.15E-01 | 1.00E+00 | 6.77E-01 |
| Biotin Metabolism | 8 | 1 | 2.16 | 5.00 | 5.25E-01 | 1.00E+00 | 6.77E-01 |
| Arginine and Proline Metabolism | 53 | 6 | 3.46 | 5.00 | 5.38E-01 | 1.00E+00 | 6.77E-01 |
| Carnitine Synthesis | 22 | 2 | 1.11 | 5.00 | 8.16E-01 | 1.00E+00 | 9.52E-01 |
| Lysine Degradation | 30 | 2 | 1.10 | 5.00 | 8.17E-01 | 1.00E+00 | 9.52E-01 |
| Porphyrin Metabolism | 40 | 1 | 0.07 | 5.00 | 9.10E-01 | 1.00E+00 | 9.52E-01 |
| Bile Acid Biosynthesis | 65 | 1 | 0.07 | 5.00 | 9.10E-01 | 1.00E+00 | 9.52E-01 |
| Cysteine Metabolism | 26 | 1 | 0.04 | 5.00 | 9.27E-01 | 1.00E+00 | 9.52E-01 |
| Folate Metabolism | 29 | 1 | 0.04 | 5.00 | 9.27E-01 | 1.00E+00 | 9.52E-01 |
| Arachidonic Acid Metabolism | 69 | 1 | 0.04 | 5.00 | 9.27E-01 | 1.00E+00 | 9.52E-01 |
| Malate-Aspartate Shuttle | 10 | 2 | 0.04 | 5.00 | 9.84E-01 | 1.00E+00 | 9.84E-01 |

**Supplemental Table 2.** Results from quantitative enrichment analysis using MetaboAnalyst 5.0 with MBRole2.0 open databases, incorporating SMPDB – CE-MS targeted bulk metabolomics of 32 metabolites in brain cortex of MDM2cKO vs. control mice.

| | Total Cmpd | Hits | Statistic Q | Expected Q | Raw p | Holm p | FDR |
|---|---|---|---|---|---|---|---|
| Threonine and 2-Oxobutanoate Degradation | 20 | 1 | 95.24 | 14.29 | 3.43E-05 | 1.34E-03 | 1.34E-03 |
| Tryptophan Metabolism | 60 | 5 | 56.19 | 14.29 | 1.36E-04 | 5.18E-03 | 2.66E-03 |
| Glutamate Metabolism | 49 | 6 | 44.07 | 14.29 | 2.47E-04 | 9.14E-03 | 2.99E-03 |
| Ammonia Recycling | 32 | 6 | 51.59 | 14.29 | 3.07E-04 | 1.10E-02 | 2.99E-03 |
| Purine Metabolism | 74 | 5 | 40.89 | 14.29 | 5.39E-04 | 1.89E-02 | 4.21E-03 |
| Aspartate Metabolism | 35 | 5 | 39.71 | 14.29 | 2.93E-03 | 9.97E-02 | 1.53E-02 |
| Pyrimidine Metabolism | 59 | 1 | 79.02 | 14.29 | 3.15E-03 | 1.04E-01 | 1.53E-02 |
| Phenylacetate Metabolism | 9 | 1 | 79.02 | 14.29 | 3.15E-03 | 1.04E-01 | 1.53E-02 |
| Propanoate Metabolism | 42 | 2 | 43.82 | 14.29 | 6.34E-03 | 1.97E-01 | 2.75E-02 |
| Methylhistidine Metabolism | 4 | 1 | 72.63 | 14.29 | 7.20E-03 | 2.16E-01 | 2.81E-02 |
| Glycine and Serine Metabolism | 59 | 8 | 33.69 | 14.29 | 1.17E-02 | 3.39E-01 | 3.97E-02 |
| Beta-Alanine Metabolism | 34 | 3 | 34.74 | 14.29 | 1.22E-02 | 3.42E-01 | 3.97E-02 |
| Histidine Metabolism | 43 | 2 | 40.80 | 14.29 | 1.52E-02 | 4.12E-01 | 4.55E-02 |
| Valine, Leucine and Isoleucine Degradation | 60 | 4 | 48.27 | 14.29 | 1.63E-02 | 4.25E-01 | 4.55E-02 |
| Urea Cycle | 29 | 6 | 29.33 | 14.29 | 2.09E-02 | 5.23E-01 | 4.76E-02 |
| Phenylalanine and Tyrosine Metabolism | 28 | 3 | 36.61 | 14.29 | 2.10E-02 | 5.23E-01 | 4.76E-02 |
| Selenoamino Acid Metabolism | 28 | 1 | 61.40 | 14.29 | 2.14E-02 | 5.23E-01 | 4.76E-02 |
| Glutathione Metabolism | 21 | 3 | 36.72 | 14.29 | 2.32E-02 | 5.23E-01 | 4.76E-02 |
| Alanine Metabolism | 17 | 3 | 36.72 | 14.29 | 2.32E-02 | 5.23E-01 | 4.76E-02 |
| Amino Sugar Metabolism | 33 | 2 | 44.00 | 14.29 | 3.40E-02 | 6.80E-01 | 6.03E-02 |
| Nicotinate and Nicotinamide Metabolism | 37 | 2 | 44.00 | 14.29 | 3.40E-02 | 6.80E-01 | 6.03E-02 |
| Warburg Effect | 58 | 2 | 44.00 | 14.29 | 3.40E-02 | 6.80E-01 | 6.03E-02 |
| Glucose-Alanine Cycle | 13 | 2 | 35.19 | 14.29 | 5.03E-02 | 8.55E-01 | 8.53E-02 |
| Methionine Metabolism | 43 | 3 | 33.31 | 14.29 | 7.45E-02 | 1.00E+00 | 1.18E-01 |
| Arginine and Proline Metabolism | 53 | 6 | 26.59 | 14.29 | 7.57E-02 | 1.00E+00 | 1.18E-01 |
| Porphyrin Metabolism | 40 | 1 | 39.77 | 14.29 | 9.37E-02 | 1.00E+00 | 1.35E-01 |
| Bile Acid Biosynthesis | 65 | 1 | 39.77 | 14.29 | 9.37E-02 | 1.00E+00 | 1.35E-01 |
| Spermidine and Spermine Biosynthesis | 18 | 2 | 30.59 | 14.29 | 9.83E-02 | 1.00E+00 | 1.35E-01 |
| Betaine Metabolism | 21 | 2 | 30.08 | 14.29 | 1.00E-01 | 1.00E+00 | 1.35E-01 |
| Carnitine Synthesis | 22 | 2 | 20.09 | 14.29 | 2.52E-01 | 1.00E+00 | 3.27E-01 |
| Malate-Aspartate Shuttle | 10 | 2 | 15.80 | 14.29 | 3.86E-01 | 1.00E+00 | 4.60E-01 |
| Catecholamine Biosynthesis | 20 | 1 | 12.56 | 14.29 | 3.89E-01 | 1.00E+00 | 4.60E-01 |
| Thyroid hormone synthesis | 13 | 1 | 12.56 | 14.29 | 3.89E-01 | 1.00E+00 | 4.60E-01 |
| Tyrosine Metabolism | 72 | 3 | 14.72 | 14.29 | 4.20E-01 | 1.00E+00 | 4.82E-01 |
| Cysteine Metabolism | 26 | 1 | 8.97 | 14.29 | 4.71E-01 | 1.00E+00 | 4.96E-01 |
| Folate Metabolism | 29 | 1 | 8.97 | 14.29 | 4.71E-01 | 1.00E+00 | 4.96E-01 |
| Arachidonic Acid Metabolism | 69 | 1 | 8.97 | 14.29 | 4.71E-01 | 1.00E+00 | 4.96E-01 |
| Lysine Degradation | 30 | 2 | 4.69 | 14.29 | 7.77E-01 | 1.00E+00 | 7.98E-01 |
| Biotin Metabolism | 8 | 1 | 0.40 | 14.29 | 8.82E-01 | 1.00E+00 | 8.82E-01 |

**Supplemental Table 3.** qPCR primers list

| Gene | Forward sequence | Reverse sequence | Reference or Catalog number |
|---|---|---|---|
| Mouse_*Gapdh* (IDT*) | TGTGTCCGTCGTGGATCTGA | CCTGCTTCACCACCTTCTTGAT | PMID: 29619376 |
| Mouse_*Cxcl1*(IDT) | TCTCCGTTACTTGGGGAC | CCACACTCAAGAATGGTCGC | PMID: 31416832 |
| Mouse_*Tnf* (IDT) | CGTCAGCCGATTTGCTATCT | CGGACTCCGCAAAGTCTAAG | PMID: 21557302. |
| Mouse_*Nox2* (IDT) | CCCTTTGGTACAGCCAGTGAAGAT | CAATCCCGGCTCCCACTAACATCA | PMID: 33907371 |
| Mouse_*Mdm2* (IDT) | GGAGTCCCGAGTTTCTCTGTG | CTTGCTGACTTACAGCCACTAAA | Detect exons 5,6 of the MDM2 gene |
| Mouse_*Hif1a* | GAA CAT CAA GTC AGC AAC GTG | TTT GAC GGA TGA GGA ATG GG | PMID: 25909079 |
| Mouse_*Cdkn1a* | NM_007669 RT² qPCR Primer Assay for Mouse Cdkn1a, GeneGlobe ID: PPM02901B-200 | | Qiagen Cat#330001 |
| Mouse_*Ccl2* | NM_011333 RT² qPCR Primer Assay for Mouse Ccl2, GeneGlobe ID - PPM03151G-200 | | Qiagen Cat# 330001 |
| Mouse_*Il1b* | NM_008361 RT² qPCR Primer Assay for Mouse Il1b, GeneGlobe ID - PPM03109F-200 | | Qiagen Cat#330001 |
| Mouse_*Il6* | NM_001314054 RT² qPCR Primer Assay for Mouse Il6, GeneGlobe ID - PPM03015A-200 | | Qiagen Cat#330001 |
| Mouse_*Trp53* | NM_022654 RT2 PCR Primer Set for Mouse Trp53tg5, Gene ID: PPM32034A-200 | | Qiagen Cat#330001 |

*IDT: Integrated DNA Technologies; Glyceraldehyde-3-phosphate dehydrogenase (*Gapdh)*

**Supplemental Table 4.** Primers list for mouse genotyping

| **Primers** | **Sequence** |
|---|---|
| FM-A | 5'-TGTGGAGAAACAGTTACTTC |
| FM-B | 5'- CTGTGCTCCTTCACAGAG |
| FM-C | 5'TGAGATGAGTCAAAGCCTGG |
| Pax8-rtTA-F | CCATGTCTAGACTGGACAAGA |
| Pax8-rtTA-R | CTCCAGGCCACATATGATTAG |
| LC1/Cre-F | TCGCTGCATTACCGGTCGATGC |
| LC1/Cre-R | CCATGAGTGAACGAACCTGGTCG |

**Supplemental Table 5.** Stages of chronic kidney disease based on eGFR categories (mL/min/1.73m$^2$) description and range*

| **Stage** | **Description** | **eGFR** |
|---|---|---|
| G1 | Normal or high | ≥90 |
| G2 | Mildly decreased | 60-89 |
| G3a | Mild to moderately decreased | 45-59 |
| G3b | Moderately to severely decreased | 30-44 |
| G4 | Severely decreased | 15-29 |
| G5 | Kidney failure | <15 |

* Reference: Levin, Adeera, et al. "Kidney Disease: Improving Global Outcomes (KDIGO) CKD Work Group. KDIGO 2012 clinical practice guideline for the evaluation and management of chronic kidney disease." Kidney international supplements 3.1 (2013): 1-150

**Supplemental Table 6.** Characteristics of the study participants, n = 26

| **Variables (mean ± SD or N (%))** | **All (n=26)** | **CKD 3b-4 (n=18)** | **CKD 5 (n=8)** |
|---|---|---|---|
| **Age, yr** | 57.08 ± 10.21 | 57.06 ± 9.08 | 57.13 ± 13.12 |
| **Female, n** | 8 (30.77%) | 5 (27.78%) | 3 (37.5%) |
| Body mass index, ($kg/m^2$) | 34.66 ± 7.72 | 35.32 ± 9.13 | 33.18 ± 2.58 |
| Systolic blood pressure, mmHg | 147.35 ± 19.35 | 144.22 ± 19.59 | 154.38 ± 17.98 |
| Diastolic blood pressure, mmHg | 75.73 ± 13.04 | 76.50 ± 12.28 | 74.00 ± 15.36 |
| **Serum creatinine, mg/dL** | 3.01 ± 1.70 | 2.03 ± 0.6 | 5.22 ± 1.19$ |
| **eGFR, mL/min/1.73m²** | 28.04 ± 13.60 | 35.83 ± 7.73 | 10.5 ± 2.67# |
| Blood hemoglobin, g/dL | 11.25 ± 2.15 | 11.79 ± 1.84 | 10.05 ± 2.41 |
| Glycated hemoglobin, % | 7.1 ± 1.1 | 7.5 ± 0.8 | 6.2 ± 1.4* |
| Urine albumin creatinine ratio, mg/g | 3715.65 ± 3038.78 | 2462.39 ± 2604.92 | 6535.50 ± 1836.72 |
| eGFR, estimated glomerular filtration rate | | | |

Two-tailed t-test, significance noted with *p<0.05 $^{\$}p < 1x10^{-6}$ ; $^{\#}p<1x10^{-8}$

**Supplemental Table 7**. Plasma-free levels of tryptophan and selective tryptophan metabolites and ratios

| **Metabolites (mean ± SD)** | All (n=26) | **CKD 3b-4 (n=18)** | **CKD 5 (n=8)** |
|---|---|---|---|
| **Tryptophan, µm/L** | 71.56 ± 22.34 | 81.04 ± 19.98 | 50.22 ± 7.86 *** |
| **Kynurenine, µm/L** | 7.21 ± 2.01 | 7.28 ± 2.08 | 7.06 ± 1.95 |
| **Kynurenic acid, µm/L** | 0.36 ± 0.20 | 0.29 ± 0.16 | 0.52 ± 0.21 ** |
| **3-Hydroxykynurenine, µm/L** | 0.03 ± 0.04 | 0.02 ± 0.03 | 0.04 ± 0.06 |
| **3-Hydroxyanthranilic acid, µm/L** | 6.20 ± 8.64 | 6.30 ± 10.17 | 5.98 ± 3.93 |
| **Quinolinic acid, µm/L** | 1.97 ± 2.42 | 1.13 ± 0.95 | 3.87 ± 3.56 ** |
| **Kynurenine/Tryptophan** | 0.11 ± 0.04 | 0.09 ± 0.03 | 0.14 ± 0.02 ** |
| **3Hydroxykynurenine/Kynurenine** | 0.003 ± 0.004 | 0.001 ± 0.002 | 0.005 ± 0.005 * |
| **Quinolinic acid/Tryptophan** | 0.03 ± 0.04 | 0.015 ± 0.013 | 0.074 ± 0.057 *** |

Two-tailed t-test, significance noted with * p < 0.05; ** p < 0.01; *** p < 0.001

**Supplemental Table 8**. Self-reported scores of brief fatigue inventory (BFI)

| **Questionnaire (mean ± SD)** | All (n=26) | **CKD 3b-4 (n=18)** | **CKD 5 (n=8)** |
|---|---|---|---|
| **BFI (total score)** | 5.15 ± 3.08 | 4.79 ± 2.99 | 5.69 ± 3.36 |

**Supplemental Methods**

**Primary renal tubular epithelial cells (pTECs):** The isolation of pTECs was performed using *Pax8*-rtTAcre;*Mdm2*f/f or *Mdm2*f/f control mice (8-10 weeks old) following established protocols (1) with slight modifications as previously described (2). After cervical dislocation, both kidneys were extracted, and the cortical regions were finely chopped and washed with PBS containing 10% Penicillin-Streptomycin (Thermo Fisher Scientific). Collagenase IV (Sigma) treatment was applied for 30 minutes at 37°C with gentle shaking. The digested tissue was then filtered through a 70 µm filter and resuspended in renal epithelial growth medium (REGM). The suspension was centrifuged at 50 g for 5 minutes at room temperature (RT), and the resulting pellet was resuspended in 5 ml of REGM. After centrifugation of the supernatant at 50 g for 5 minutes at RT, the second pellet containing tubular cells was resuspended in 5 ml of REGM. Both pellet suspensions were combined (total of 20 ml) and centrifuged again at 50 g for 5 minutes at RT. The final pellet was resuspended in 2 ml of REGM and plated onto a collagen coated T25 flask. Cells were cultured under 5% CO2 at 37°C, with media changed every 48 hours. After 8-10 days, tubular cells were passaged for further experiments, and all studies were performed using passage 1 cells. pTECs isolated from *Pax8*-rtTAcre;*Mdm2*f/f or *Mdm2*f/f control mice were treated for 48 h with 1µg/ml doxycycline to induce *Mdm2* depletion (3).

**Protein extraction and western blot:** Protein extraction from brain tissue (RIPA buffer) was followed by quantification using BCA assay (Thermofisher #23235), and equal protein quantities were loaded (12 µl). Transfer on PVDF membranes was followed by blocking (LICOR #927-60001) and incubation with primary (Mouse-anti-MDM2 #PIMA124643; 1:500 or Beta-Actin β-Actin Antibody #4967; 1:5000) and secondary antibodies (Goat anti-Rabbit IgG #926-32211 and goat anti-mouse IgG #926-68070; 1:10,000). Protein bands were visualized using LICOR Odysse CLx.
